# Phylogenetic Novelty Scores: a New Approach for Weighting Genetic Sequences

**DOI:** 10.1101/2020.12.03.410100

**Authors:** Nicola De Maio, Alexander V. Alekseyenko, William J. Coleman-Smith, Fabio Pardi, Marc A. Suchard, Asif U. Tamuri, Jakub Truszkowski, Nick Goldman

## Abstract

**Background:** Many important applications in bioinformatics, including sequence alignment and protein family profiling, employ sequence weighting schemes to mitigate the effects of non-independence of homologous sequences and under- or over-representation of certain taxa in a dataset. These schemes aim to assign high weights to sequences that are ‘novel’ compared to the others in the same dataset, and low weights to sequences that are over-represented.

**Results:** We formalise this principle by rigorously defining the evolutionary ‘novelty’ of a sequence within an alignment. This results in new sequence weights that we call ‘phylogenetic novelty scores’. These scores have various desirable properties, and we showcase their use by considering, as an example application, the inference of character frequencies at an alignment column — important, for example, in protein family profiling. We give computationally efficient algorithms for calculating our scores and, using simulations, show that they improve the accuracy of character frequency estimation compared to existing sequence weighting schemes.

**Conclusions:** Our phylogenetic novelty scores can be useful when an evolutionarily meaningful system for adjusting for uneven taxon sampling is desired. They have numerous possible applications, including estimation of evolutionary conservation scores and sequence logos, identification of targets in conservation biology, and improving and measuring sequence alignment accuracy.

## Background

Some of the most popular applications in bioinformatics, including multiple sequence alignment [1], sequence database search [2] and protein family profiling [3, 4], employ sequence weighting schemes as a way to mitigate the effects of non-independence of homologous sequences. For example, a database may contain many closely related sequences from one species (like humans) and its close relatives, while other more distantly related species might be under-represented. To address possible problems associated with these biases, several sequence weighting schemes have been proposed over the years. These methods assign a score to each sequence considered, with the aim of assigning reduced weights to sequences from over-represented clades and larger weights to sequences from under-represented clades.

PSI-BLAST [2], for example, employs the Henikoff and Henikoff (1994) weighting scheme [5] where the score of a sequence is the average of the scores of each position of the sequence, the score of a position being 1*/rd*, with *r* the number of different characters at the considered alignment column and *d* the number of times the character of the considered sequence and position appears in the considered alignment column. The idea of this weighting scheme is to give equal weight to all characters observed at one alignment column, dividing this weight equally among those sequences sharing that character at that position. This method has the advantage of being very fast to calculate, and of giving higher weights to sequences with more rare characters that are, therefore, likely more distantly related.

HMMER [6] and the CLUSTAL family of aligners [1, 7, 8] use the weighting scheme of Gerstein et al. [9] (similar to [10]), which defines sequence weights iteratively along a phylogeny from tips to root. At each step, the length of the considered tree branch is split proportionally to the current weights of its descendant sequences, and is then added to the weights of the descendant sequences. Here, the idea is that weights are determined by divergence between groups of sequences. The more diverged one group of sequences is from the others, the higher weights it will have. However, the weight of a group is shared among the sequences in the group, so that in a group with many similar sequences each of those sequences will have small individual weight.

Other sequence weighting schemes have also been proposed, although they have seen fewer applications. Maximum discrimination sequence weighting [11] is a complex approach that aims to optimally distinguish homology from chance alignments in database searches. Henikoff and Henikoff (1992) proposed a method that splits sequences into clusters based on sequence similarity, and assigns equal weights to sequences in the same cluster and a total weight of 1 to each cluster [12]. Vingron and Argos weighted sequences proportionally to their average distances from all other sequences [13]; Sibbald and Argos proposed an approach in which a sequence receives more weight if it is more isolated in sequence space [14]. Altschul et al. measured evolutionary correlations among sequences using branch lengths in the phylogeny, and then calculated sequence weights using the inverse of the variance-covariance matrix [15]; Gotoh developed a fast approximation of this method [16]. Similar ideas have also been explored within methods aimed at estimating character frequencies at a given position in a protein [17, 18, 19], defining tree or alignment informativeness [20, 21, 22, 23, 24], and quantifying diversity within a habitat and prioritising conservation efforts [25, 26, 27, 28, 29, 30, 31].

The many weighting schemes proposed have rarely been assessed and compared under different scenarios. We show that these approaches tend to suffer from limitations, for example requiring an ultrametric or rooted phylogenetic tree, or being unable to cope with certain levels of sequence divergence (e.g. identical sequences or very diverged sequences). Instead, here we propose a new weighting scheme, the first to be derived from the idea of evolutionary ‘novelty’. We quantify the novelty of each sequence compared to the other sequences under consideration by computing the probability that, at a given position, sequences are ‘phylogenetically identical by descent’ (PIBD): that is, that they descended from a common ancestor without any substitution occurring. This rigorous approach allows us to quantify the novelty of sequences in very general scenarios (without specific assumptions regarding the phylogeny relating the considered sequences) while being robust to uneven sampling and very elevated or reduced divergence levels, and generally conforming to guiding principles for an acceptable weighting scheme [32]. We present algorithms and scripts to efficiently compute these weights from a phylogeny and from a multiple sequence alignment.

As shown by the examples above, this new weighting scheme has a number of possible applications, from gene family profiles and multiple sequence alignment evaluation to ecology and conservation biology. Here, we focus on the inference of sitewise character frequencies. Inference of character frequencies at a given alignment column is not only important for gene family profiling, but also for modeling evolutionary fitness, calculating conservation scores, and visualizing sequence logos [33, 34, 35, 36, 37]. We show that our methods result in efficient and accurate inference of character frequencies, with clear advantages compared to previous sequence weighting schemes.

## Methods

### Phylogenetic novelty scores

We consider a phylogenetic tree *ϕ* describing the evolutionary relationships of its *N* tips *s*_1_, …, *s*_*N*_. We want to define weights *w*_*s*_ representing how ‘novel’ tip *s* is compared to the other tips of *ϕ*. Throughout this paper we consider the tips to represent biomolecular sequences comprising amino acid or nucleotide characters, but other possible sets of characters could equally be accommodated. We assume that we have one sequence associated with each tip, conveniently sharing the same names *s*_1_, …, *s*_*N*_, and arranged as the rows of an alignment *A*. We start by defining weights that are a function of *ϕ* only, and so depend on the evolutionary history relating the considered sequences and not on the specific sequences themselves. In the next section we extend the definitions to also condition on the observed sequence characters.

As a motivating example, if *ϕ* consists of only extremely long branches, then we want *w*_1_ = … = *w*_*N*_ = 1. In fact, in this case, all sequences represent effectively independent observations, so no weighting correction is needed. This means that, unlike many sequence weighting schemes (e.g. [38, 39, 14, 9]), we want to account for the effect of saturation, so that doubling the length of a long tree branch has negligible effect on the weights.

If instead *ϕ* has branches all of length 0, we want *w*_1_ = … = *w*_*N*_ = 1*/N*, so that the total alignment score is 1, as in [38, 39, 14, 9]. This is because all the observed sequences are now just perfectly dependent copies, and so in total they represent just one independent observation of a sequence. At an intermediate level, if *ϕ* has two tips (*N* = 2), and branch length such that half of the ancestral characters are expected not to have mutated in either branch (they are PIBD with probability 0.5), then we want the total alignment score to be 1.5, and both weights to be 0.75; this is because in this case only half of each sequence will be novel with respect to the other, so in total we observe 1.5 novel sequences, and we want the two sequences to have the same weight.

A simple way to describe how novel *s*_1_ is with respect to *s*_2_ could be to count the number of mismatches between their sequences. However, even if *s*_1_ and *s*_2_ were very divergent from each other, their sequences would still be identical at some alignment column because of chance or of convergent evolution, instead of close relatedness. In our approach, *s*_1_ can be novel with respect to *s*_2_ at a column of *A* even if they share the same character, as long as they are not PIBD.

We usually cannot know for sure if sequences are PIBD and at which alignment column, so we define *p*_*s*_(*i*) as the probability that, at a generic alignment column, the number of tips of *ϕ* (including *s*) that are PIBD to *s, i*_*ϕ*_(*s*), is exactly *i*. For example, *p*_*s*_(*N*) is the probability that, at a generic alignment column, no substitution occurs along *ϕ*; *p*_*s*_(1) is the probability that no tip (except *s*) in *ϕ* is PIBD to *s* at some arbitrary alignment column. We then define the weight *w*_*s*_ of *s* within *ϕ* as:

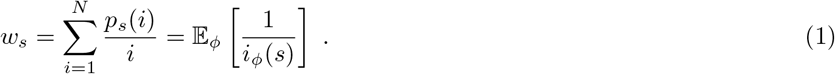

In the simplest case of nucleotide sequences evolving under the JC69 substitution model ([40]; all substitution rates are 1*/*3), the probability that two nodes in *ϕ* separated by branch length *t* are PIBD is *e*^−*t*^. So, again in the simple case that *N* = 2 and that the two branches in *ϕ* have each length *t/*2, *s*_1_ and *s*_2_ each have weight 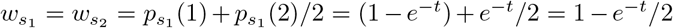, and the sum of the weights is 2 − *e*^−*t*^. The same is true for any pair of branch lengths with sum *t*, of course.

We expect the definition of sequence weights given by Equation 1 to be useful for character frequency inference and many other applications. In fact, in addition to satisfying classical sequence weighting requirements [32], these *w*_*s*_ can also be efficiently calculated from any *ϕ* and substitution model, as discussed later. We refer to weights *w*_*s*_ as the ‘phylogenetic novelty scores’ (PNS). We call the sum of all weights in *ϕ* the ‘effective sequence number’ (ESN): 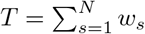, representing the expected number of evolutionarily distinct character observations at an alignment column. An example graphical representation of the PNS is shown in Figure 1

**Figure 1.**
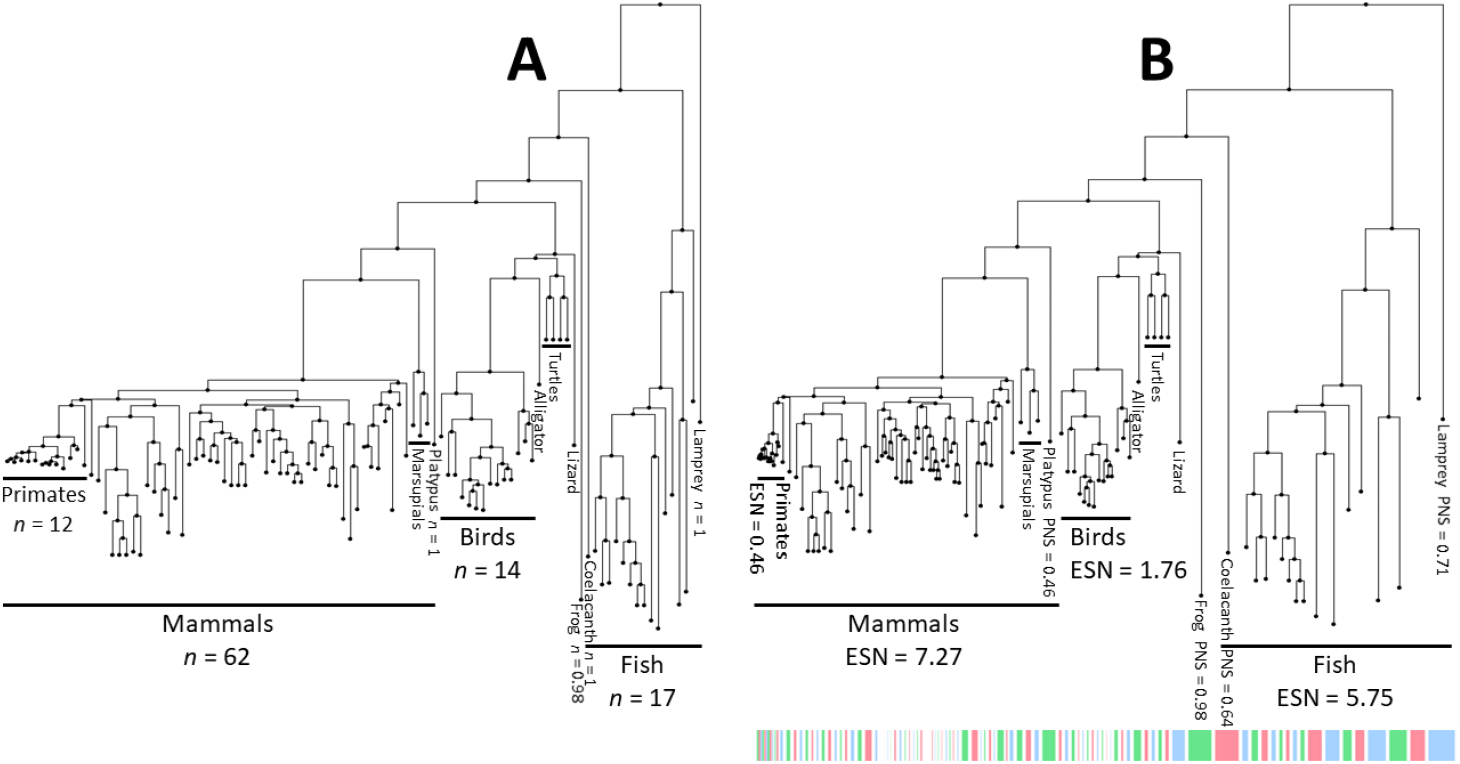
Example of PNS for a 100-vertebrates tree. Here we show graphically the values of the phylogenetic novelty scores *w*_*s*_ from Equation 1 for the tips of a tree of 100-vertebrate species. The tree is taken from the UCSC genome browser 100-way alignment of vertebrates to the human genome, downloaded from http://hgdownload.cse.ucsc.edu/goldenPath/hg38/multiz100way/hg38.100way.commonNames.nh. This tree was also used for simulations in this work. **A:** The tree has all tips spaced uniformly on the horizontal axis, representing the case of no weighting scheme being used. **B:** Tips are spaced horizontally according to their *w*_*s*_ weight. The weight of each tip can also be seen in the length of the colored bars. Notice how species in regions of the tree with many close relatives (e.g. mammal, primate and bird clades) have low PNSs, and so take up less space individually. This means the horizontal dimension of the plot now gives more equal representation of the novelty of each sequence and clade, instead of emphasising densely sampled clades. More divergent species with few close relatives (e.g. lamprey, coelacanth, frog and platypus) have higher PNSs and are given more horizontal space, representing the greater novelty of their sequences relative to other species in the tree. Cumulative ESN scores (clade-wise sum of PNSs) are also shown for some clades.

#### Conditioning on observed data

In this section we define weights that are a function not only of phylogeny *ϕ*, but also of a specific alignment column *D* of alignment *A*. These weights refer not to the novelty of a sequence *s*, but of its specific character *D*_*s*_ observed in row *s* of column *D*. The probability that two tips of *ϕ* are PIBD at a specific alignment column can be strongly affected by the observed characters at that column. Clearly, if the two tips differ at alignment column *D* then the probability that they are PIBD, conditional on *D*, is 0. The case that the two tips have the same character in *D* is less trivial. If we assume that the two tips are separated by a total divergence time *t*, and for simplicity assuming a JC69 substitution model [40], then the probability that the two tips have the same nucleotide is (1 + 3*e*^−4*t/*3^)*/*4 and the probability that the two tips are PIBD is *e*^−*t*^; therefore, the probability that the two tips are PIBD conditional on them having the same nucleotide is 4*e*^−*t*^*/*(1 + 3*e*^−4*t/*3^).

We denote by *p*_*s*_(*i*|*D*) the probability that exactly *i* sequences are PIBD to *s* at the given, observed alignment column *D*. The new positional PNSs conditional on *D* are then defined as:

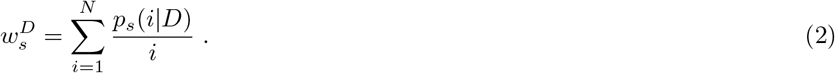

### Algorithms for Calculating the Phylogenetic Novelty Scores

We present several algorithms for calculating PNS. One of these methods (‘up-down pruning’) is the most computationally efficient, and so is described below, but may also be the most difficult to understand. For this reason, we also mention other approaches and include their full description in the Supplement. In the following we assume that the phylogeny *ϕ* is rooted; the case of an unrooted topology follows simply by placing an arbitrary root on the tree, as long as the substitution process is reversible and at equilibrium (as in this case the scores are not affected by the position of the root). In the case of non-reversible or non-stationary character evolution, the position of the root can affect the scores, and so a rooted phylogeny (which could in principle be estimated from sequences in this scenario) is required.

#### Calculating PNS scores via simulation

We can calculate PNS by simulating sequence evolution along *ϕ*. If we are interested in weights *w*_*s*_, at each iteration we start by sampling a root character from the equilibrium distribution. We then sample its descendant characters and the mutation events along the branches of *ϕ* using standard methods (e.g. [41], “method 2” of [42], [43]) until we reach all the tips, recording each substitution that occurs and hence which tip characters are PIBD. Note that it is not possible to achieve this using software such as *evolver* [44] or “method 1” of *INDELible* [42] that only simulate the start and end state of each branch, and do not distinguish between characters that are PIBD and those that happen to match following multiple substitutions. For each iteration we associate a score of 1*/i* to a tip of *ϕ* if its observed character is PIBD to the characters of exactly *i* tips. The final weight of a tip is then obtained by averaging its scores over all iterations. Weights 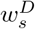 can be similarly calculated employing a variant of the up-down approach [45] to sample characters at internal nodes of the phylogeny conditional on *D*. A straightforward but inefficient way to achieve the same result is to simulate characters without any conditioning, and discard those iterations that do not match *D*. These approaches are described in more detail in the Supplement.

#### Calculation of PNS scores via brute-force

We can calculate PNS via brute-force, that is, by enumerating all possible mutational histories on *ϕ* by considering all possible character assignments at each end of each branch, and for each branch considering whether there is at least one substitution on it or not. Each mutational history results in a score as in the previous method using simulation. By averaging the scores of all mutational histories, while accounting for different probabilities of different histories, we can calculate the *w*_*s*_ or 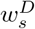 weights. The full methods are described in detail in the Supplement.

#### Pruning method to calculate the ESN

The ESNs 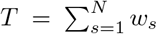 or 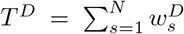 can be calculated very efficiently (computational cost 𝒪(*N*)) if one is not interested in the weights of the individual tips. The idea is to calculate *T* iteratively on each subtree of *ϕ* starting from the tips until we reach the root. We call this the ‘pruning ESN’ method, due to its similarity with Felsenstein’s pruning algorithm [46]. See the Supplement for details.

#### Up-down pruning approach to calculate PNS

We now present the main, efficient algorithm that we use and recommend for calculating PNS. It can be used for calculating either *w*_*s*_ or 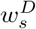 weights and can be considered an adaptation of Felsenstein’s pruning algorithm [46]. The method visits all nodes in *ϕ* starting from the tips and toward the root (‘up’ phase) and then again a second time starting from the root and moving downward to the tips (‘down’ phase), similar to the up-down approach of [45]. The computational cost of this algorithm is cubic in the number of tips *N*.

In the following we assume that the substitution rate matrix *Q* is given. The probability of having character *k* at the end of a branch of length *t*, conditional on having character *j* at its start, is then 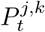, the entry in row *j* and column *k* of *P*_*t*_ = exp(*tQ*). See [47] for a more detailed introduction to these concepts in molecular phylogenetics. Starting with character *j* at the top node of a branch of length *t*, we denote the probability that no substitution occurs along the branch, and therefore that the top and bottom nodes of the branch are PIBD, as:

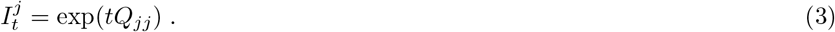

Our objective is to calculate, for each tip *s* of *ϕ*, the probability distribution (*p*_*s*_(0) = 0, *p*_*s*_(1), …, *p*_*s*_(*N*)) of having each possible number of PIBD sequences (defined as in Equations 1 and 2). To address both the cases of 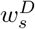 and *w*_*s*_, we present our description as conditioned on data *D*; for the case that one is interested in *w*_*s*_, the same equations can be used but setting *D* as non-informative. (For example, with DNA sequences a non-informative column *D* will have all entries equal to character “N”, representing an unknown nucleotide, so that the partial likelihood for column *D* at each tip is 1.) As before, we denote the observed character at tip *s* by *D*_*s*_; we now represent the observed characters for the leaves in sub-phylogeny *ϕ*′ of *ϕ* as *D*_*ϕ′*_. In the particular case that *ϕ*′ = *ϕ*, we have *D*_*ϕ*_ = *D*, so we can represent the final values of interest for tip *s* also as *p*_*s*_(*i*|*D*_*ϕ*_).

For most of the following, we condition probabilities on information from only part of *ϕ*. Given a node *v* of *ϕ*, and given a sub-phylogeny *ϕ*′ of *ϕ*, we define 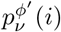 to be the probability that there are exactly *i* tips in *ϕ*′ that are PIBD to *v*. We also define 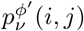 as the probability of having *i* tips in *ϕ*′ that are PIBD to *v* and to have character *j* in *v*. Similarly, we define 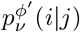 to be the probability of having *i* tips in *ϕ* that are PIBD to *v*, conditional on having character *j* in *v*. Finally, 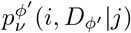 is the probability that *i* tips in *ϕ*′ are PIBD to *v* and that the observed data in *ϕ*′ is *D*_*ϕ′*_, conditional on having character *j* in *v*.

The first step of the up phase is to initialise 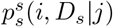 at every tip *s* of *ϕ*, for every character *j*, and for 0 :⩽ *i* :⩽ *N* :

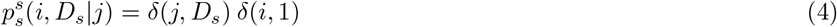

where *δ*(*x, y*) is the Kronecker delta function (*δ*(*x, y*) = 1 if and only if *x* = *y*; *δ*(*x, y*) = 0 otherwise).

Next, starting from the tips, we move iteratively ‘upward’, toward the root of *ϕ*. If branch *b* with length *t* connects the two nodes *v*_1_ (the parent or upper node) and *v*_2_ (child or lower node), then *b* splits *ϕ* into two sub-phylogenies. We call these *ϕ*_1_ and *ϕ*_2_, with *ϕ*_2_ the sub-phylogeny of *ϕ* containing *v*_2_ (but not *b*) and all its descendant nodes and branches, and *ϕ*_1_ the sub-phylogeny of *ϕ* containing all nodes and branches (except *b*) not in *ϕ*_2_. Assuming that we have already visited all branches and nodes below *b*, and therefore know 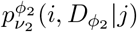 for every character *j* and every 0 :⩽ *i* :⩽ *N*, we can then calculate 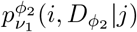 for every character *j* and every 0 :⩽ *i* :⩽ *N* :

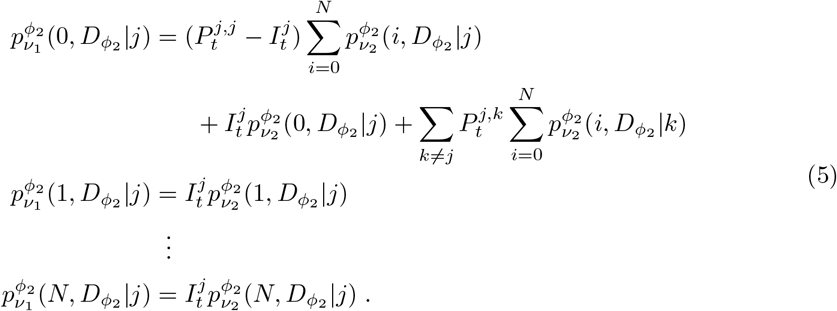

Many of the 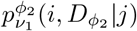 will be 0 (when *i* is larger than the number of tips in *ϕ*_2_). In practice, we have made use of this to speed up the implementation of the algorithm, but we ignore it here for brevity.

Thanks to Equation 5 we can ‘move’ probabilities up along branches, starting from the initialisations at the tips. Next, we show how to ‘merge’ probabilities when we reach an internal node *v*. A given internal node *v* splits *ϕ* into three sub-phylogenies (a parent one, *ϕ*_*P*_, a left child one *ϕ*_*L*_, and a right child one *ϕ*_*R*_), each associated with one of the three branches adjacent to *v* (one parent and two child branches). If *v* is the root, then for simplicity we consider its parent sub-phylogeny to exist but be empty. Assuming that we have already visited all branches and nodes descendant of *v*, and therefore know 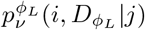 and 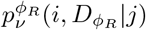 for every character *j* and every 0 :⩽ *i* :⩽ *N*, and denoting by *ϕ*_*L*_ ∪ *ϕ*_*R*_ the sub-phylogeny obtained by joining sub-phylogenies *ϕ*_*L*_ and *ϕ*_*R*_, we can calculate 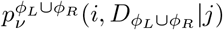 for every character *j* and every 0 :⩽ *i* :⩽ *N* :

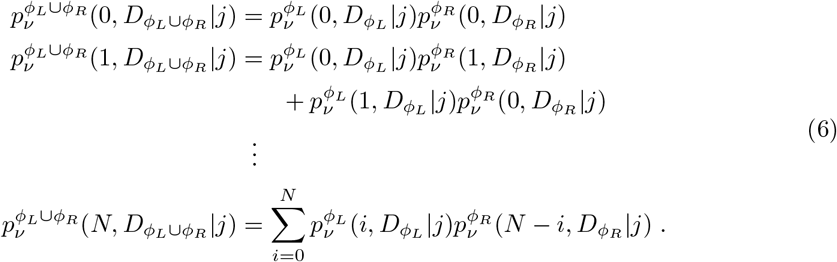

Equation 6 is one of the most computationally demanding steps of the algorithm (jointly with Equation 10 below) as it has up to quadratic cost in *N*. Equation 6 is used on each internal node of *ϕ*, and so causes the algorithm to have a total time complexity in the order of 𝒪 (*N* ^3^).

Using Equations 5 and 6 iteratively, we can calculate 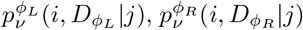 and 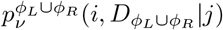 for each internal node *v*, each 0 :⩽ *i* :⩽ *N* and any character *j*. We stop once we reach node *ρ*, the root of *ϕ*. At *ρ* we have 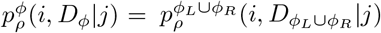 for any character *j* and 0 :⩽ *i* :⩽ *N*. If *π* are the character frequencies at *ρ*, we then have the joint probabilities:

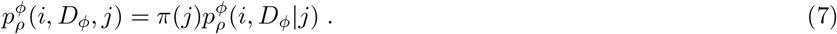

This concludes the ‘up’ stage of the method, which is more succinctly described in Equation 8:

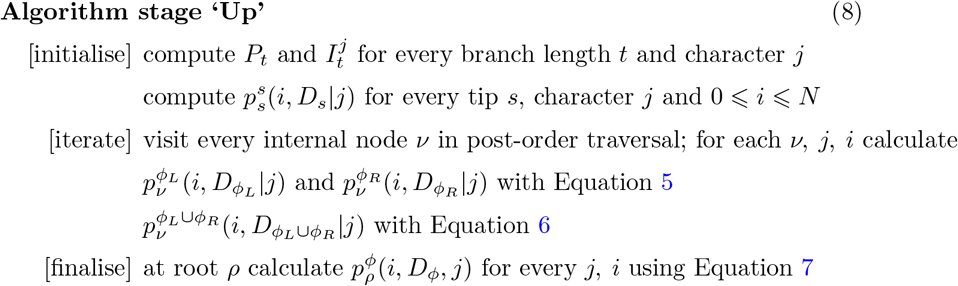

The ‘down’ phase is the second and last stage of the algorithm. Starting from root *ρ*, we move toward the tips, visiting each node and branch in pre-order traversal. Given branch *b* of length *t* connecting nodes *v*_1_ (parent) and *v*_2_ (child), we assume, as in Equation 5, that *ϕ*_1_ and *ϕ*_2_ are the two sub-phylogenies induced by *b*. Assuming that we have already visited iteratively all ancestor branches of *b*, and therefore know 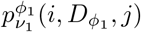 for every character *j* and 0 :⩽ *i* :⩽ *N*, we can calculate 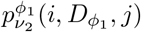 for every character *j* and 0 :⩽ *i* :⩽ *N* :

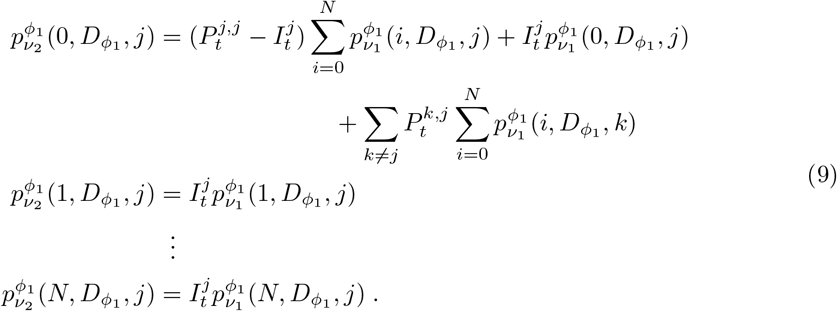

Equation 9 allows us to ‘move’ probabilities downward along branches, starting from the root. Next, we show again how to ‘merge’ probabilities when we reach an internal node *v*. Given one left child sub-phylogeny *ϕ*_*L*_ of *v*, and given its parent sub-phylogeny *ϕ*_*P*_, we can calculate 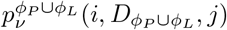 for every character *j* and 0 :⩽ *i* :⩽ *N*:

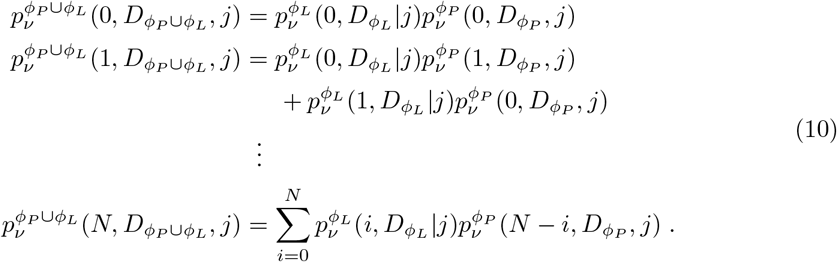

We use Equation 10 twice for each internal node *v*, once with the left child sub-phylogeny *ϕ*_*L*_ and once replacing *ϕ*_*L*_ with the right child sub-phylogeny *ϕ*_*R*_. Using Equations 9 and 10 iteratively, we calculate 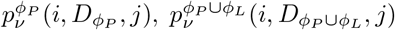 and 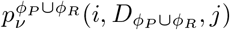 for each internal node *v*, each 0 :⩽ *i* :⩽ *N* and every character *j*. After we have visited every internal node of *ϕ*, we reach all tips *s* using Equation 9 to obtain 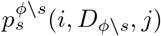 for all characters *j* and all 0 :⩽ *i* :⩽ *N*, where *ϕ* \ *s* is the sub-phylogeny obtained by removing *s* (and its parent branch) from *ϕ*. We then combine these probabilities at the tips with the initialisation probabilities 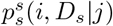 to obtain, at every tip *s*, for all characters *j* and each 1 :⩽ *i* :⩽ *N* :

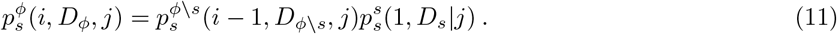

The final probabilities of interest, 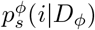, can be calculated for every 0 :⩽ *i* :⩽ *N* and every tip *s* as:

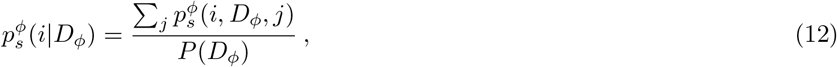

where *P* (*D*_*ϕ*_) is the probability of the data (the phylogenetic likelihood of *ϕ* and the substitution model for *D*). In the case *D* is empty, that is, if we want to calculate weights *w*_*s*_, then *P* (*D*_*ϕ*_) = 1. Otherwise, if we are interested in weights 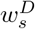, then *P* (*D*_*ϕ*_) can be calculated as a normalisation factor such that the probabilities 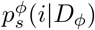 at any *s* sum over *i* to 1. In either case, the final PNS scores can be easily calculated substituting the results from Equation 12 into Equation 1 or 2.

We summarise the ‘down’ stage of the algorithm in Equation 13:

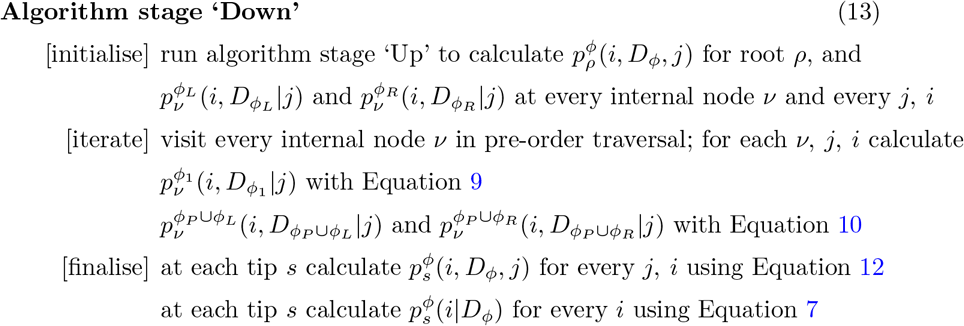

### Fast approximation

The most efficient algorithm above has cubic cost in *N*. In some circumstances, for example when *N >* 10^5^, it becomes important to consider faster solutions. For this reason, we also present an approximate PNS that can be calculated more efficiently. With *i*_*ϕ*_(*s*) the random variable representing the number of tips in *ϕ* that are PIBD to *s*, we have 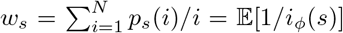 (Equation 1). As an alternative fast approximation we consider:

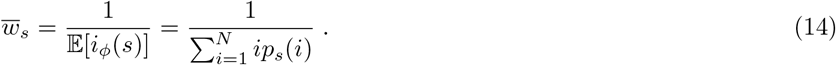

The weights 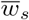 can be computed very efficiently with an up-down pruning approach, requiring only 𝒪(*N*) time, so we refer to them as ‘fast PNS’. The algorithm to calculate weights 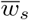 has many similarities to the one in the previous Section, and is described in detail in the Supplement.

### Application to inference of character frequencies

Inference of character frequencies specifically for a single alignment column has broad applications such as modeling selection [33, 34], and creating profile HMMs [3, 4, 6] and sequence logos [35, 36, 37]. Here, we assume that the frequencies of interest are the equilibrium frequencies at a given alignment column, i.e. the average character frequencies over long evolutionary times. Such frequencies are typically represented in molecular phylogenetics as *π*, with *π*(*j*) being the equilibrium distribution of character *j* [47]. This definition of frequencies fits well with the assumptions of profile HMMs, and is also reasonable for sequence logos, although we acknowledge that different definitions might be also considered in different settings. In this work, we want to investigate and compare different methods for inferring *π*.

The simplest inference method is to use the observed frequency *p*(*j*) of character *j* within the given column as an estimate of the true frequency *π*(*j*). This approach corresponds to assuming that all sequences are independent of each other. This approach might be ideal in some circumstances, for example when the considered sequences are not homologous but only evolutionary convergent, but might be inappropriate in others. As an example, consider an alignment of 1000 homologous human sequences and two mouse sequences (1002 homologous sequences in total). Genetic variation within mice, and variation between mice and humans will have negligible effects on estimates *p*(*j*), which will be dominated by within-human genetic variation. However, human sequences are highly correlated, as they have very short divergence time between each other, so within-human allele frequencies will typically not represent evolutionary equilibrium character distributions. The problem here is that using *p*(*j*) as an estimate of *π*(*j*) means treating homologous sequences as independent of each other, while they are often strongly correlated due to shared evolutionary histories.

A traditional way to address this problem is to use sequence weights, for example our *w*_*s*_, to reduce the contribution of groups of closely related sequences. We can in fact define *p*^*w*^(*j*), a new estimate of *π*(*j*), as:

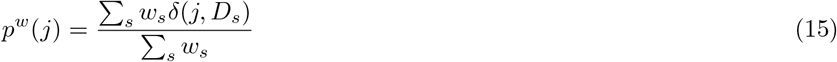

where, as before, *D*_*s*_ is the observed character for sequence *s* at the alignment column *D* under consideration, and *δ* is again the Kronecker delta function.

We investigate and compare the performance, for character frequency inference, of the three weighting schemes introduced above: 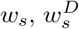 and 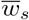. We also consider two popular sequence weights: those defined by [5], which we call HH94, and by [9], which we call GSC94. HH94 first calculates, for any *s*, the score of *s* at an alignment column *D*, which we will denote 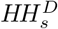. This score 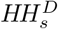 is 1*/rd*, with *r* the number of different characters in *D* and *d* the number of times character *D*_*s*_ appears in *D*. The weight for sequence *s*, which we denote *HH*_*s*_, is then defined as the average of 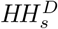 over all columns *D* of alignment *A*.

GSC94 defines sequence weights iteratively along a phylogeny, by visiting branches in post-order traversal (from the tips to the root). First, all terminal branches (those connected to the tips of *ϕ*) are visited, and the length of a terminal branch connected to tip *s* is assigned as the initialisation value of the weight *GSC*_*s*_ of *s*. Then, every time an internal branch *b* is visited, its length *t* is distributed among the weights of its descendant sequences. More precisely, first *t* is split among the tips, with the part *t*_*s*_ assigned to *s* being 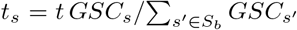, where *S*_*b*_ is the set of tips descendent from *b*. Secondly, each *GSC*_*s*_ for *s* ∈ *S*_*b*_ is increased by *t*_*s*_. After the last branches connected to the root have been processed, the *GSC*_*s*_ are the final GSC94 weights.

In addition to the character frequency inference *p*(*j*) based on the observed frequencies, and *p*^*w*^(*j*) based on one of many weighting schemes studied, we also consider character frequency inference via phylogenetic maximum likelihood (ML). We perform this using PhyML v3.1 by fixing the phylogenetic tree (inferred from the whole alignment *A* using FastTree v2.1.10 [48]) and the substitution model exchangeabilities, and inferring, one alignment column *D* at a time, only the equilibrium character frequencies *π*.

#### Bayesian approaches to character frequency inference

Above, we introduced point estimate methods for character frequency inference. These methods do not measure inference uncertainty, and this can result in a very limited summary of the available data. For example, observing character *j* 100 times in an alignment column from 100 distantly related species leads all above methods to infer 100% frequency for *j*; however, so also does observing *j* two times within an alignment column of just two closely related sequences. While in the first scenario there should be little uncertainty regarding the inferred frequencies, in the second scenario uncertainty should be elevated. Using a Bayesian method is a natural way to address this issue, and also allows the inclusion of priors over characters frequencies. Here we present a Bayesian variant of the weight-based character frequency inference of Equation 15.

If *A* is composed of *N* independent (non-homologous) sequences, the likelihood of a column *D* is *P*(*D*|*π*) = Π_*s*_*π* (*D*_*s*_).. It is simple to combine this likelihood with a character frequency prior to obtain a Bayesian posterior distribution, and perform Bayesian character frequency inference. However, we are interested in the general case where the sequences in *A* are related by a phylogeny *ϕ*, and therefore are not independent. One possible way to perform Bayesian inference of *π* in this scenario would be using Bayesian phylogenetic methods such as BEAST [49] or MRBAYES [50], but at often excessive computational cost. Instead, we propose an approximation of the likelihood function *P* (*D*|*π*) based on sequence weights *w*_*s*_:

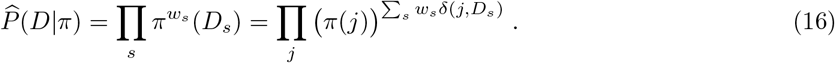

Similarly, we can replace weights *w*_*s*_ in Equation 16 with any other weighting scheme. In the following, we assume a uniform prior *P* (*π*) on character frequencies, meaning that all possible *π* are similarly likely a *priori*. Alternative priors are possible, and some might be more realistic, but usually at the cost of introducing more parameters in the model. Our approximation of the posterior probability *P* (*π*|*D*) is then:

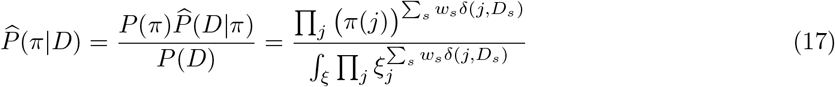

where the integral in the denominator is over all possible character frequencies *ξ*. Equation 17 is a Dirichlet distribution with parameters *α*_*j*_ = 1 + Σ_*s*_*w*_*s*_*δ*(*j, D*_*s*_), so in the following we use the properties of Dirichlet distributions [51]. The (approximate) maximum *a posteriori* and ML *π* are both given by the weighted observed character frequencies *p*^*w*^(*j*) in Equation 15. The approximation of the expectation of *π*(*j*), *E*(*π*(*j*)|*D*) is however:

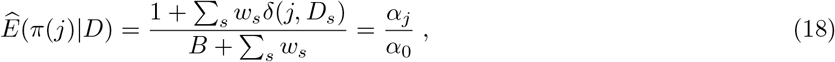

where *B* is the number of possible characters in the considered alphabet, *α*_*j*_ = 1 + Σ_*s*_*w*_*s*_*δ*(*j, D*_0_) and *α*_*j*_ = *B* + Σ_*s*_*w*_*s*_. This can be seen as the ML estimate of *π* in the presence of 1 pseudo-count per character. The posterior variance is then approximated as:

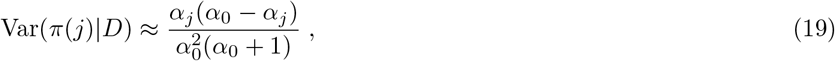

which can be used as a measure of the uncertainty over character frequencies. However, considering that Equation 17 has beta-distributed univariate marginals 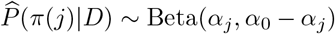, in the following we derive approximate 95% posterior probability intervals using the stats.beta.ppf function in *scipy* [52].

### Simulations

We use simulations to test and compare computational demands of calculating PNS values as well as for assessing the accuracy of different approaches to infer position-specific character frequencies. In the base simulation scenario, we simulate nucleotide sequence evolution along a 100 vertebrate taxa phylogeny (Figure 1) using Dendropy [53]. We use a HKY85 substitution model [54] with transition:transversion ratio *κ* = 3 both for simulation and inference. We simulate 10 replicates, each replicate consisting of an alignment of 1000 columns. Alignment columns are evolved independently of each other (conditional on the tree and the substitution model).

Specific equilibrium character frequencies *π* are assigned to each alignment column. For each alignment, 800 columns (80%) are simulated as evolving under the same background equilibrium character frequency distribution, which we set to *π*(*A*) = *π*(*T*) = 0.3 and *π*(*C*) = *π*(*G*) = 0.2. The background character frequency distribution represents, in our simulations, the evolutionary dynamics of positions not strongly affected by selective forces; at these positions, the equilibrium character frequency distribution is constant because it is mostly determined by neutral mutational biases, which we assume constant across all alignment columns. The remaining 20% of alignment columns are simulated under position-specific selection, with position-specific equilibrium character frequency *π* sampled from a Dirichlet distribution prior with *α* = 0.1 (Supplementary Figure S1A).

Our aim is, for each replicate and each alignment column, to infer *π* from the simulated sequences alone. For each replicate/alignment, we first infer a phylogenetic tree and alignment-wide HKY85 substitution model parameters using FastTree v2.1.10 [48]. We then consider this tree and the HKY85 *κ* parameter to be fixed and infer column-specific character equilibrium frequencies. While *π* is inferred separately at each column, the HKY85 alignment-wide parameters (including nucleotide frequencies) inferred with FastTree are used in some sequence weighting schemes (for example, in Equation 3). The methods we used to infer equilibrium frequencies are:

- the observed character frequencies in the alignment column (the *p*(*j*) described above),
- observed frequencies corrected using the HH94 [5] weights and Equation 15,
- observed frequencies corrected using the GSC94 [9] weights and Equation 15,
- observed frequencies corrected using the PNS weights *w*_*s*_ from Equation 1 combined with Equation 15,
- observed frequencies corrected using our PNS weights conditional on data, 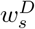 from Equation 2, combined with Equation 15,
- observed frequencies corrected using our fast approximate PNS 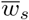 weights (Equation 14) combined with Equation 15,
- Bayesian variants of the methods above, and
- ML phylogenetic inference (only of equilibrium character frequencies) with PhyML v3.1 [55].

All the methods above, except FastTree and PhyML, were implemented in custom Python scripts available from https://bitbucket.org/nicofmay/noveltyscores.

In addition to the basic simulation scenario, we also consider variant scenarios in order to investigate how certain parameters can affect the results:

- We consider alternative tree lengths, which we obtain by multiplying all branch lengths in the tree in Figure 1 by a constant coefficient, either 0.2 or 5.
- We consider the case of amino acid characters instead of nucleotides. In this case, we simulate under an LG substitution model [56], and when we do inference we assume that the substitution model (including character frequencies) is known. Column-specific character frequencies are inferred as usual. In this case, equilibrium character frequencies for columns under selection are sampled from a Dirichlet distribution prior with *α* = 0.02 (Supplementary Figure S1B).
- To test the effect of very biased taxon sampling in an alignment, we added multiple (either 100 or 1000) human tips to the tree in Figure 1. The short phylogeny relating the human sequences was randomly sampled at each replicate under a standard coalescent prior [57] with mean coalescent time 0.001 between human sequences. This short human phylogeny was then appended to the human tip of the tree in Figure 1.
- To test the robustness of methods to the assumption of an ultrametric tree (a tree where all tips have equal distance from the root), we consider the case of a strongly non-ultrametric trees, as is common for some viruses such as influenza.

## Results

### Sequence weights

We implemented all the considered weighting schemes and all simulations within custom Python scripts (https://bitbucket.org/nicofmay/noveltyscores), making use of the phylogenetic python package dendropy [53]. We implemented all the algorithms presented in the Methods section for calculating weights *w*_*s*_ and 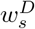, and used comparisons of weights from different algorithms to assess the correctness of the implementations.

PNS shows similar trends to previous weighting schemes HH94 and GSC94, assigning higher weights to phylogenetically isolated taxa and smaller weights to taxa within clades with many other closely related taxa (Figure 2). In particular, PNS seems to show an intermediate ‘intensity’ compared to the two other schemes. GSC94 weights are the most extreme, assigning the highest weights of any scheme to the most evolutionarily isolated taxa in the tree of Figure 1, such as Lamprey, Coelacanth and frog *Xenopus tropicalis*. For example, for Lamprey, the GSC94 normalized weight is about 3 times larger than HH94, while PNS is about 2 times larger than HH94. Conversely, for taxa in over-represented clades, such as Human, GSC94 gives the smallest weight, HH94’s weight being many times larger, and PNS being intermediate. Rescaling the branch lengths of the tree does not change this overall trend (Supplementary Figure S2).

**Figure 2.**
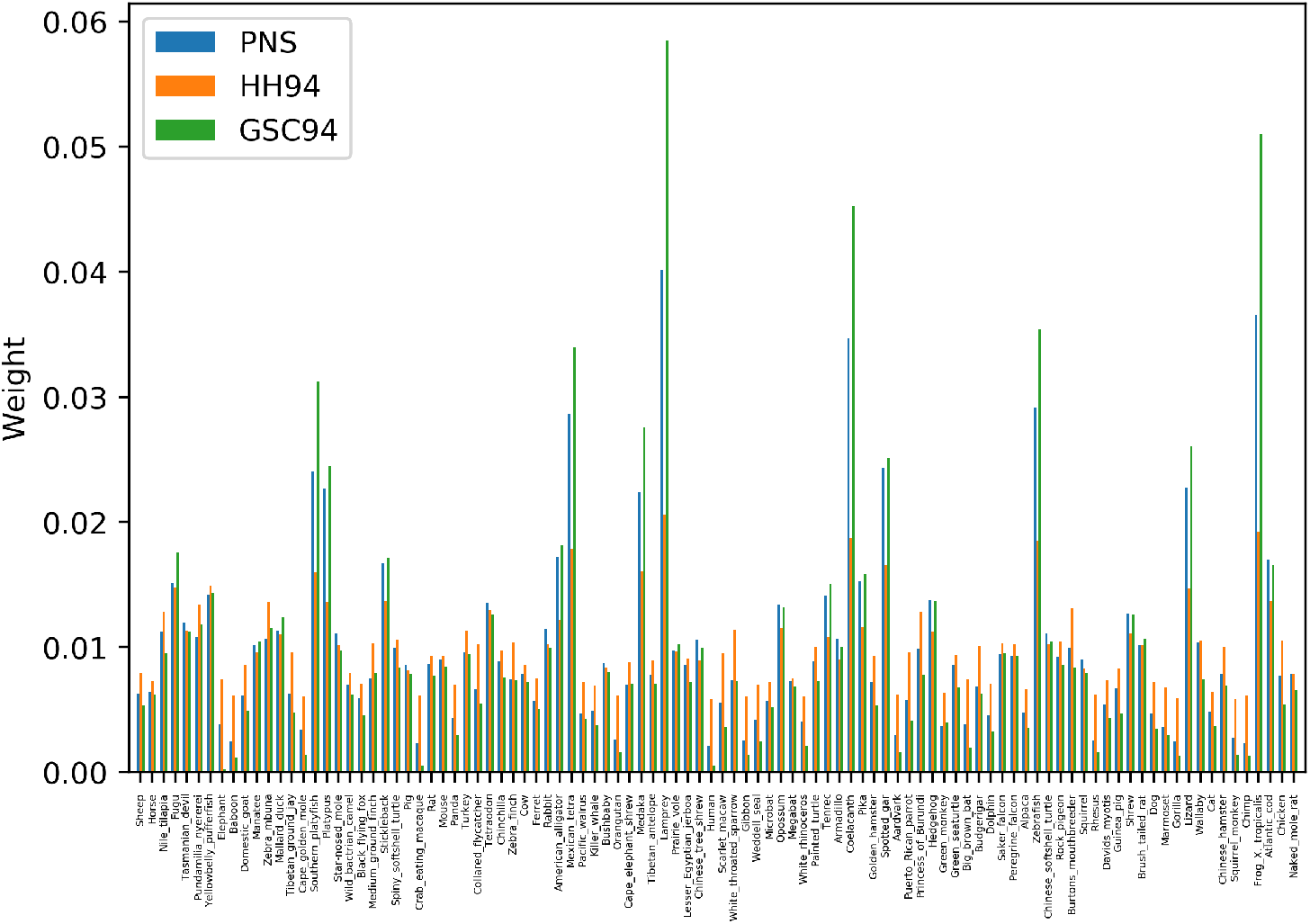
Comparison of different weighting schemes. Bars show weights assigned to the tips of tree in Figure 1 (species names on *x*-axis labels) in the scenario of nucleotide data (1 locus of 1kb) by different weighting schemes: PNS (weights *w*_*s*_), HH94 [5] and GSC94 [9]. Weights from each scheme are normalized so that the sum over taxa is 1.

### Computational demand

Calculating sequence weights based on a phylogeny usually requires limited computational demand, with the dominant cost being the estimation of the phylogeny (Figure 3 and Supplementary Figure S3). One exception to this are the 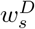 weights of Equation 2: these, being conditional on the data observed at a specific alignment column, need to be re-computed for each position. Calculation of these weights requires time cubic in *N*, and so it is not surprising that these weights are slower than phylogenetic inference. The other slowest method for character frequency is phylogenetic ML (PhyML), which also needs to be run once for each alignment column. All other approaches are at least one order of magnitude faster than PhyML and 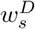 in estimating character frequencies, and are practical also for much larger trees (e.g. thousands of taxa, see Figure S3). Calculating weights 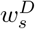 and estimating frequencies by ML, instead, becomes infeasible on such larger trees. Estimating frequencies using HH94 weights is the fastest of the methods considered, as it does not require prior estimation of a phylogenetic tree. The second fastest is GSC94, followed by the 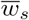 weights, and finally by the *w*_*s*_ weights.

**Figure 3.**
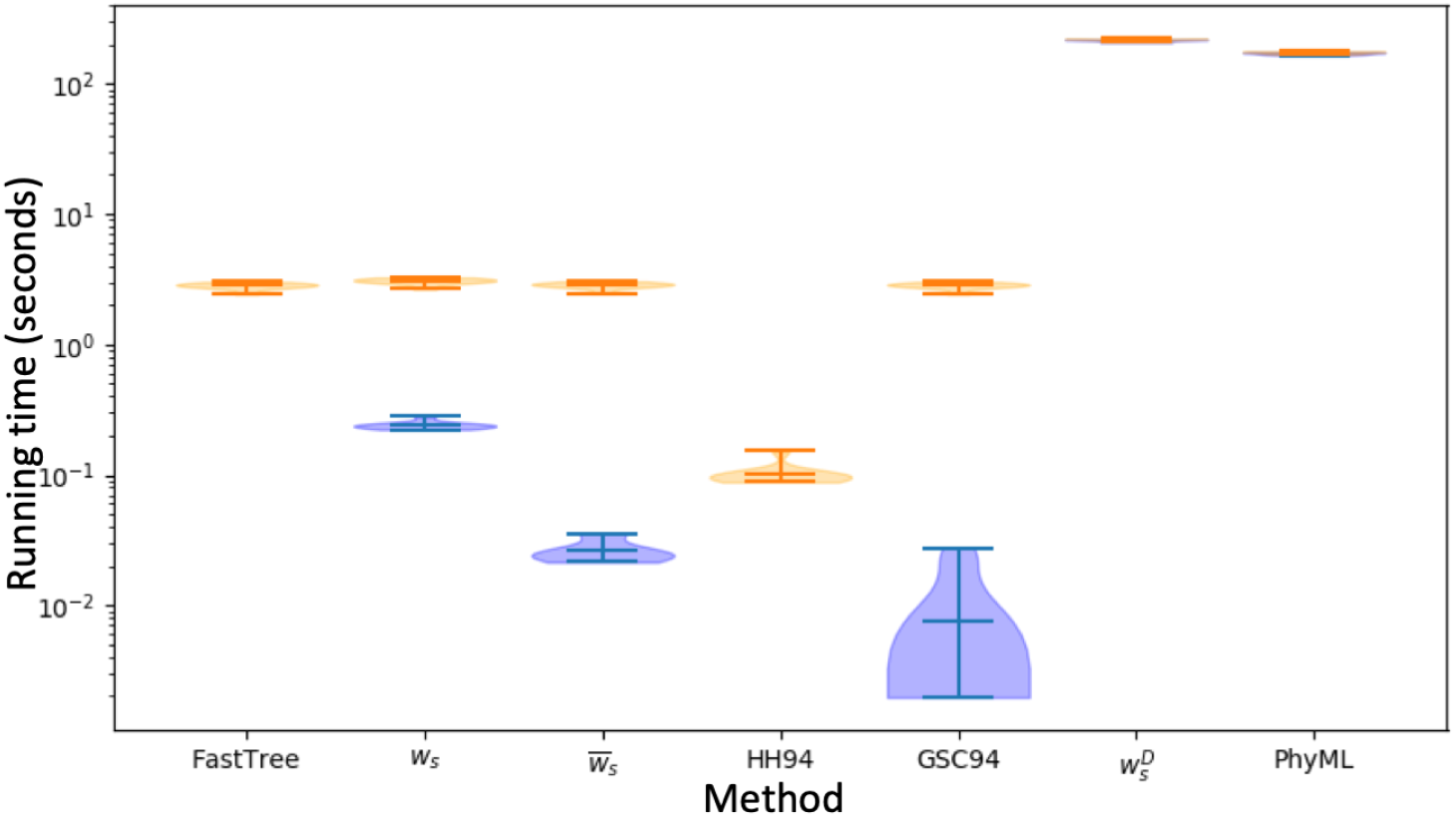
Computational demand of different approaches to character frequency estimation. Violin plots summarise the running times, in seconds, of different methods. All analyses were run on a MacBook Pro 2017. Each plot contains values for 10 replicates of the scenario of the unscaled tree in Figure 1 and nucleotide data. Time cost for computing frequencies from un-weigthed observed characters is not shown as it is negligible. Time demand of Bayesian variants of PNS weights is also not shown, as it is the same as for their non-Bayesian variants (Bayesian variants only require the addition of pseudocounts compared to non-Bayesian variants). ‘FastTree’ represents the cost of running phylogenetic inference with FastTree prior to weight calculation. Orange violin plots show the total cost (including computational cost of phylogenetic inference for methods requiring a phylogeny). Blue violin plots show the cost of calculating the scores without taking into account the cost of phylogenetic tree inference. For 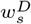 and ‘PhyML’, blue and orange plots overlap. Calculating HH94 weights is, overall, the fastest approach among those considered here, as it does not require phylogenetic inference.

### Accuracy of character frequency inference

Here we assess the ability of different weighting schemes, including those derived from our new PNS methods, to facilitate inference of column-specific character frequencies. We measure of the accuracy of an approach by calculating, at each alignment column, the Euclidean distance between simulated and inferred character frequencies. ML phylogenetic inference with PhyML is almost always the most accurate method (Figures 4 and 5). This is perhaps not surprising, given that this approach fully models the effects of varying equilibrium character distributions on character evolution along the phylogeny. However, this approach is also the most computationally demanding, and the advantage of schemes based on sequence weights is that they can be much faster, in particular on datasets with many sequences or many alignment columns. The only case where PhyML seems marginally less accurate than weights-based methods is at high divergence and strong selection (Figure 5F). This is probably due to the particular implementation in PhyML, which does not allow character frequencies below 1%.

**Figure 4.**
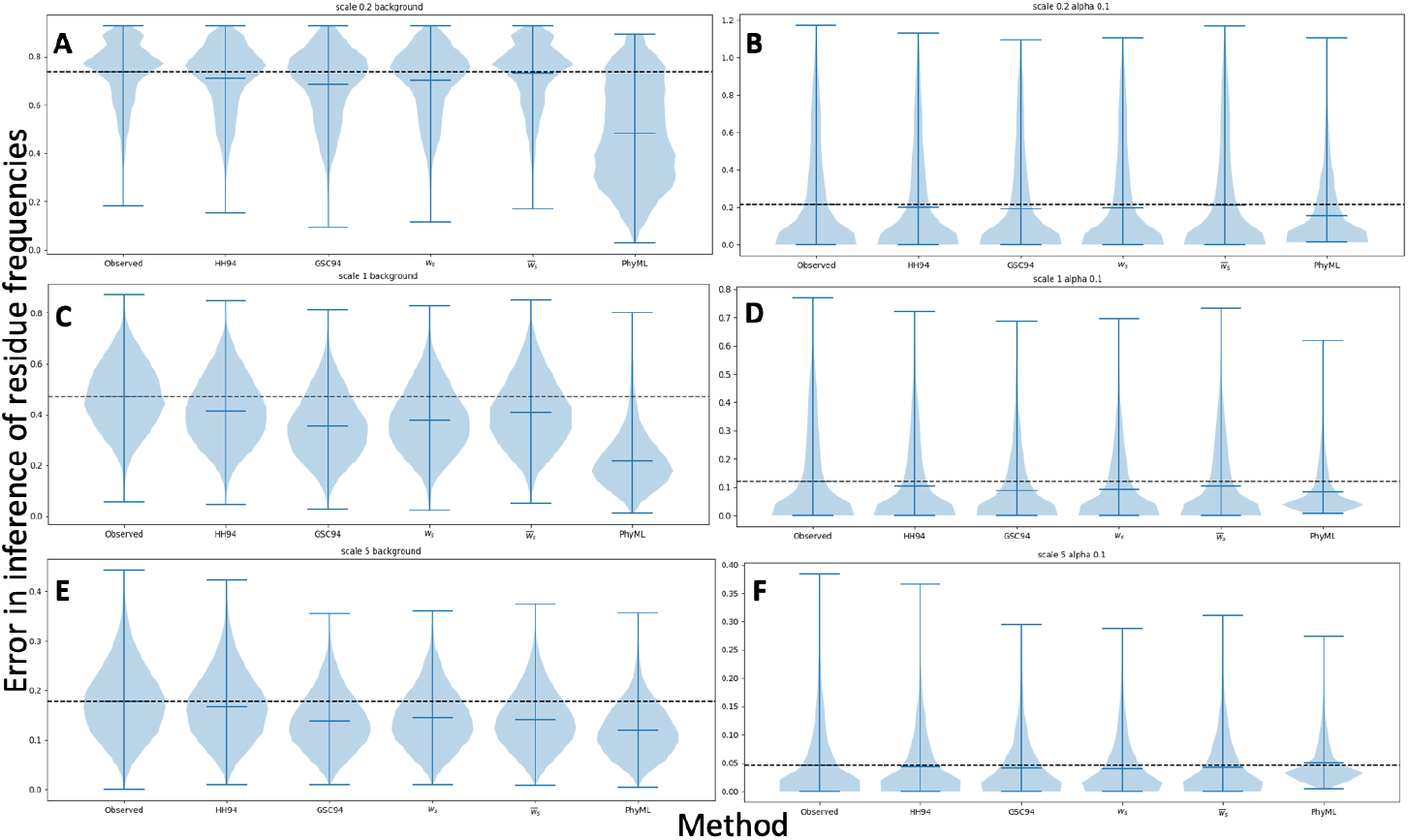
Equilibrium frequency inference error. Comparison of the accuracy of different methods for reconstructing equilibrium frequencies in the basic simulation scenario (nucleotide characters and tree as in Figure 1). Violin plots summarise the nucleotide frequency inference error (on the *y*-axis), measured as the Euclidean distance between the vectors of column-specific simulated nucleotide frequencies and inferred ones. Each plot contains 10 replicates, and each replicate contains 800 alignment columns evolved under the background nucleotide frequencies (**A, C** and **E**), or 200 alignment columns evolved under equilibrium nucleotide frequencies sampled from a Dirichlet distribution with *α* = 0.1 (**B, D** and **F**). Horizontal black dashed lines aid comparison by showing the median error of the first method (frequencies extracted from character counts). In **A** and **B** the tree branch lengths were scaled by a factor of 0.2; in **C** and **D** by a factor of 1.0; and in **E** and **F** by a factor of 5.0. Each plot shows results for a particular character frequency inference method, indicated on the *x*-axis. Results from additional methods (e.g. Bayesian approaches) are shown in Supplementary Figure S4.

**Figure 5.**
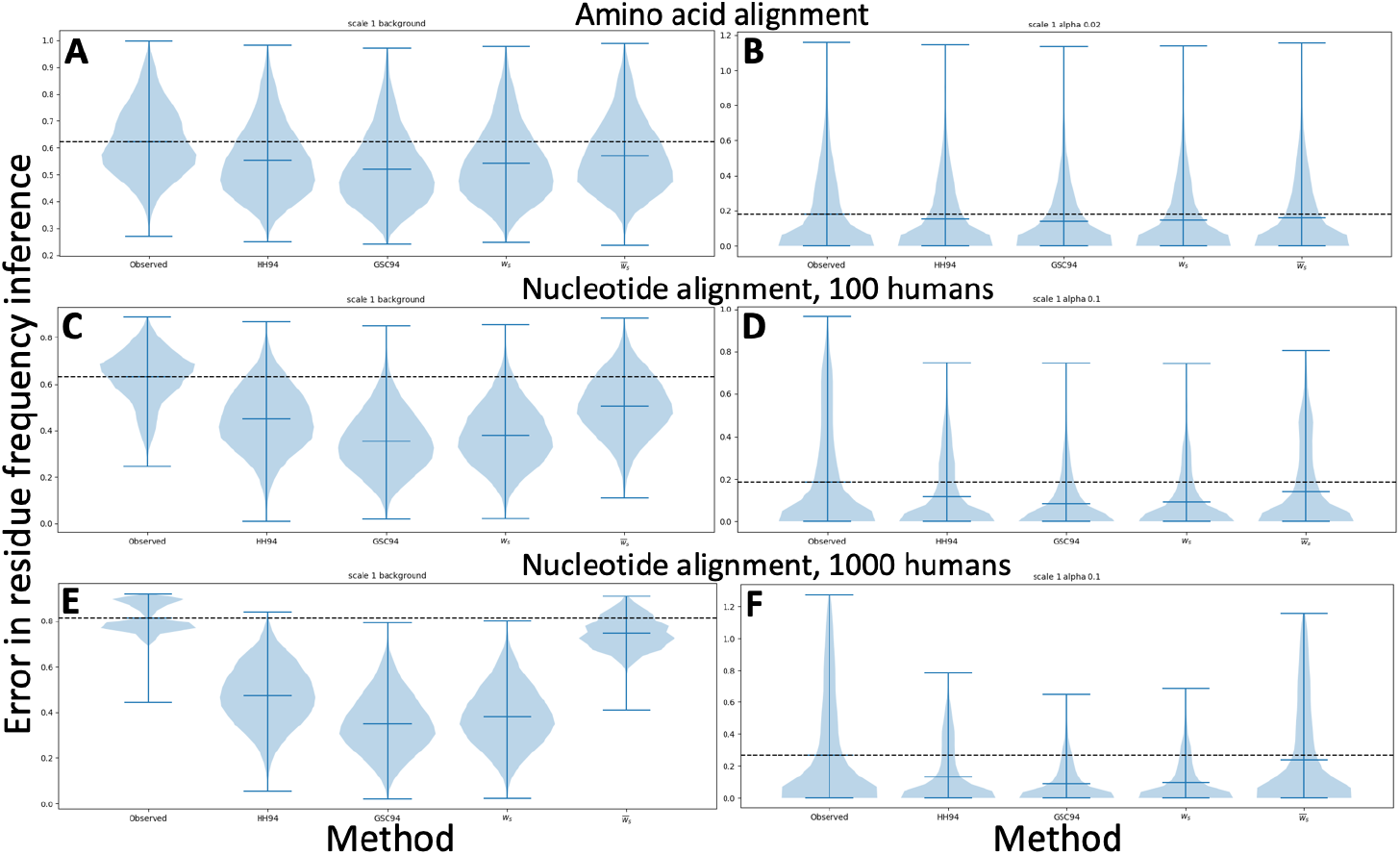
Equilibrium frequency inference error under different scenarios. Similarly to Figure 4, we compare the accuracy of different methods for reconstructing equilibrium frequencies. However, here we consider the simulation scenarios of amino acid sequences and modified trees with increased over-representation of human sequences. Values shown are as in Figure 4. Each plot contains 10 replicates, and each replicate contains 800 alignment columns evolved under the background character frequencies (**A, C** and **E**), or 200 alignment columns evolved under equilibrium character frequencies sampled from a Dirichlet distribution with *α* = 0.1 for **D** and **F** and *α* = 0.02 for **B**. In **A** and **B** simulations are under the tree in Figure 1 and with amino acid sequences. In **C** and **D** we consider nucleotide sequences and the tree in Figure 1 with 100 added human sequences (see Methods). In **E** and **F** we instead add 1000 human sequences. Results from additional methods (e.g. Bayesian approaches) are shown in Supplementary Figure S4. Results from PhyML are not available, due to excessive computational demand.

All the weighting schemes considered improve character frequency inference compared to the simplest approach of counting the observed characters in a alignment column (Figure 4 and 5). GSC94 and *w*_*s*_ weights seem to give more accurate results than HH94 and 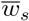 weights, in particular within very biased datasets (Figure 5**C-F**). The latter is not too surprising, given that weights 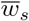 are an approximation of weights *w*_*s*_.

We note that in Figures 4 and 5 the weights *w*_*s*_ and GSC94 give very similar accuracy, with GSC94 sometimes marginally outperforming *w*_*s*_. In theory, we expect the weights *w*_*s*_, compared to GSC94, to benefit from the advantages of being based on intuitive mathematical principles and accounting for the effects of saturation (which however might be limited in many scenarios). A further limitation of the GSC94 weights is that they do not work well with trees in which tips have very different distances from root (non-ultrametric trees). This effect has limited impact in our basic simulation scenario, as the tree in Figure 1 is not far from ultrametric. However, when we consider a strongly non-ultrametric tree (Figure 6**A**), as is often observed for some viruses such as influenza [58, 59], we see that the GSC94 weights are strongly impacted, resulting in considerably worse inference than any of the other weighting schemes studied, and worse even than observed character frequencies (Figure 6**B**). The reason is that, in such strongly non-ultrametric trees, GSC94 weights at terminal, younger tips tend to be considerably larger than GSC94 weigths at older tips closer to internal nodes and in particular those closer to the root. Even in cases when observed characters close to internal nodes can provide useful information regarding equilibrium frequencies, for example when branches are sufficiently long in Figure 6A, GSC94 weights are still almost exclusively distributed on the latest two phylogenetic tips in this scenario.

**Figure 6.**
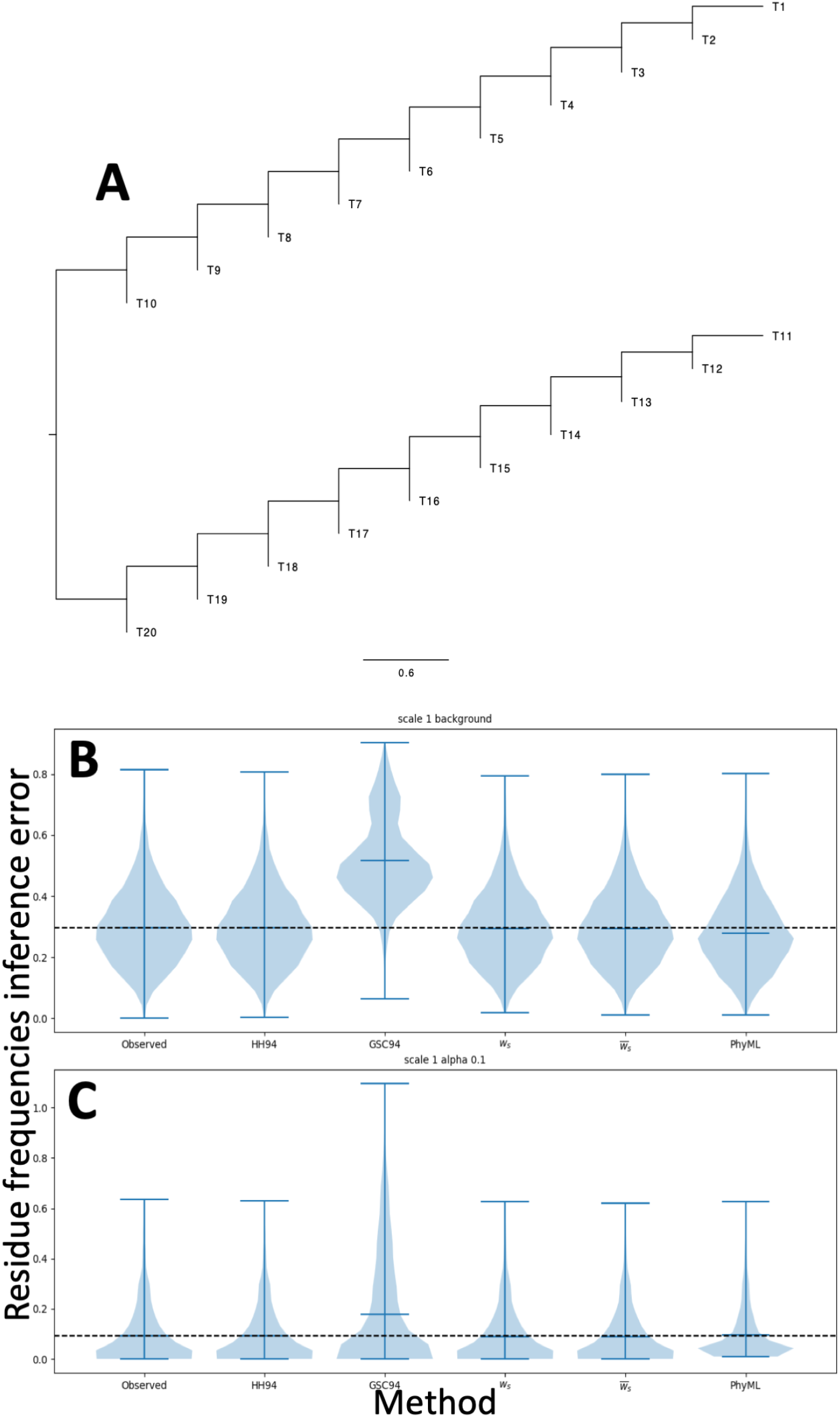
Equilibrium frequency inference error with a strongly non-ultrametric tree. **A**: The strongly non-ultrametric phylogenetic tree under which simulations for this figure are performed. Some tips of the tree (e.g. T10, T20) are close to the root while others (T1, T11) are considerably more evolutionarily distant; in an ultrametric tree, all tips would instead have the same distance from the root. **B** and **C**: Violin plots summarising nucleotide frequency inference error (*y*-axis), measured as the Euclidean distance between the vectors of column-specific simulated nucleotide frequencies and inferred ones. Each plot contains 10 replicates, and each replicate contains (**B**) 800 alignment columns evolved under the background nucleotide frequencies, or (**C**) 200 alignment columns evolved under equilibrium nucleotide frequencies sampled from a Dirichlet distribution with *α* = 0.1. Each plot refers to a particular character frequency inference method, indicated on the *x*-axis.

Using sequence weights conditional on the data at the specific column, i.e. 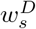 from Equation 2, does not seem to improve accuracy (Supplementary Figure S4) while, as shown in Figure 3, it does significantly impact computational demand. Using a Bayesian approach to character frequency inference means that the prior on character frequencies can affect the result of the inference. This can have a positive effect if the prior distribution is based on reliable evidence from sources other than the currently considered dataset. However, in our simulations we consider a completely arbitrary prior (corresponding to observing one character of each type at the considered alignment column) and this has the effect of slightly shifting the inferred frequencies closer to a uniform distribution (Supplementary Figure S4). Expectedly, this overall improves character inference at sites evolving under the background frequencies, while it worsens inference at sites evolving under strong selection.

## Discussion

We have proposed a new approach for assigning weights to the sequences in an alignment, or, equivalently, to the tips of a phylogenetic tree. First, we define phylogenetic novelty scores (PNS) based on rigorous mathematical principles. These scores summarise how novel is a sequence (respectively, tip), in evolutionary terms, with respect to the rest of the alignment (respectively, tree) and have a number of desirable properties, including meeting the objective criteria of [32].

We have showcased our scores’ potential use by considering, as an example application, the inference of position-specific character frequencies. We demonstrate, using simulations, that our scores improve accuracy of character frequency estimation compared to some popular sequence weighting schemes such as HH94 [5] and GSC94 [9]. While the improvement over HH94 weights is evident in most simulations, PNS and GSC94 weights show very similar performance. However, we demonstrate that, unlike GSC94 weights, PNS are not affected when the assumption of tree ultrametricity is violated. Over most scenarios, the best method for position-specific character frequency inference seems to be standard phylogenetic

ML inference; however, this approach is also very computationally demanding, and is not suitable for large datasets.

Character frequency inference, our example use-case for PNS, has a number of important applications. Character frequencies are fundamental parameters used in HMM profiling of protein families [3, 6], and our scores could therefore improve approaches to this task. Our scores could also be used to improve character frequency estimates used within alignment column-wise conservation scores [35, 36, 37], frequently defined as

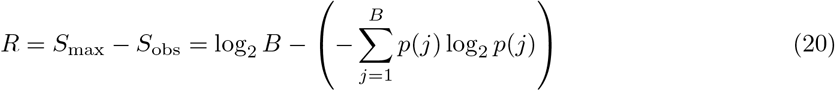

where *p*(*j*) is the frequency of character *j* at a given alignment column, and *B* is the number of characters (*B* = 4 for nucleotides and *B* = 20 for amino acids). (*S*_max_ is the maximum possible entropy at the considered position, equal to log2 *B*, while *S*_obs_ is the observed value.) Typically, the *p*(*j*) are inferred from the observed character frequencies at an alignment column; however, as we have shown, our PNS can significantly improve the inference of these frequencies, and therefore of conservation scores. Our simulations suggest that this is in fact the case (Supplementary Figure S5).

Sequence weights, like our PNS, also have many other applications, for example to aid alignment inference. They have been shown to improve sequence alignment [1] and profile searches [10, 5], and examples of their use include PSI-BLAST [2] (which uses HH94 weights) and the CLUSTAL family of aligners (e.g. [10, 1] use GSC94 weights). Our scores could therefore result in improved alignments.

Sequence weights are also used to measure alignment quality, and our scores could be used for example in the context of the information content score (ICS) [60] or the norMD approaches [61]. Furthermore, our scores could be used in measures of conservation priority in conservation biology, such as phylogenetic diversity *PD* [25], quadratic diversity *Q* [30] and the phylogenetic entropy index *H*_P_ [31].

Lastly, we note that our scores could be used to improve the definition of phylogenetic effective sample size to be used for AICc [62] and BIC [63]. This is usually defined as the number of alignment columns, but this is not the only reasonable choice [64, 65].

## Conclusions

We have proposed a new type of sequence weights that benefit from a number of favourable properties and are derived from rigorous mathematical evolutionary principles. When inferring character frequencies, we showed that these sequence weights offer considerable computational advantage over full phylogenetic ML estimation, and are more accurate than alternative sequence weighting schemes. Thanks to their computational efficiency and robustness to phylogenetic assumptions, our phylogenetic novelty scores could have a positive impact in a number of fields, from sequence alignment and protein family profiling to phylogenetics and conservation biology.

## List of abbreviations

PIBD: ‘phylogenetically identical by descent’
PNS: ‘phylogenetic novelty scores’
ESN: ‘effective sequence number’
HH94: weighting scheme by Henikoff and Henikoff, 1994
GSC94: weighting scheme by Gerstein, Sonnhammer and Chothia, 1994
HKY85: substitution model by Hasegawa, Kishino and Yano 1985
LG: substitution model by Le and Gascuel, 2008

## Authors’ contributions

NG devised the concept of Phylogenetic Novelty Scores (Equation 1) as a weighting scheme for evolutionarily related organisms. FP performed early computations of the fast approximation (Equation 14). AVA and MAS contributed ideas toward the pruning methods described here. AUT performed simulations to derive estimates of *ws* and 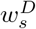 (the latter both ‘inefficiently’ and efficiently!). JT created an early implementation of the up-down pruning algorithm. WJC-S created code for plotting phylogenies scaled by PNS values (Figure 1). NDM combined all these elements, added others, completed all theory and algorithms, implemented the methods, and ran the analyses presented. NDM and NG wrote the manuscript, which was read and approved by all the authors.

## Acknowledgements

We thank Julia de Beer, Sean Eddy and István Miklós for helpful discussions on these topics.

## Ethics approval and consent to participate

Not applicable.

## Consent for publication

Not applicable

## Availability of data and materials

All scripts and data are available from https://bitbucket.org/nicofmay/noveltyscores.

## Additional Files

Additional file 1 — Supplement

File containing all Supplementary Figures and additional details of the methods.

