## Supplement for "Phylogenetic Novelty Scores: a New Approach for Weighting Genetic Sequences"

Nicola De Maio[1*], Alexander V. Alekseyenko[1,2], William J. Coleman-Smith[1,3], Fabio Pardi[1,4], Marc A. Suchard[5], Asif U. Tamuri[1,6], Jakub Truszkowski[1,7] and Nick Goldman[1]

[1]European Molecular Biology Laboratory, European Bioinformatics Institute (EMBL-EBI), Wellcome Genome Campus, Hinxton, UK
[2]Current address: Department of Public Health Sciences, Medical University of South Carolina, Charleston, SC, USA
[4]Current address: LIRMM, University of Montpellier, CNRS, Montpellier, France
[5]Departments of Biostatistics, Biomathematics and Human Genetics, University of California, Los Angeles, CA, USA
[6]Current address: Research Computing, University College London, London, UK
[7]Current address: RBC Borealis AI, Waterloo, Ontario, Canada

[*]

### Supplementary Figures

**Figure S1. Simulated character frequency distributions.** Graphical representation of the Dirichlet distributions used for sampling (**A**) simulated equilibrium nucleotide frequency distributions ($\alpha = 0.1$), and (**B**) simulated equilibrium amino acid frequency distributions ($\alpha = 0.02$). Each plot refers to the probability of the frequency of one character. On the $x$-axes are ranges of character frequencies, and on the $y$-axis is the probability of such ranges under the given Dirichlet distribution.

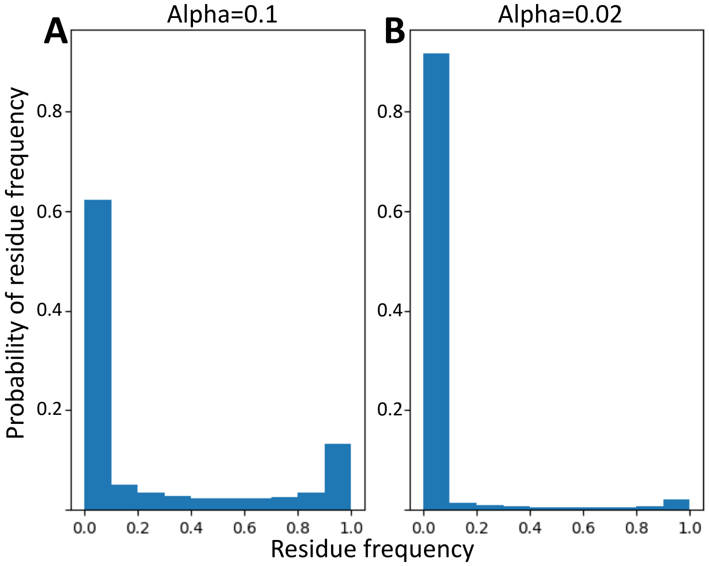


**Figure S2. Comparison of different weighting schemes with rescaled branch lengths.** Bars show weights assigned to the tips of tree in Figure 1 (species names on $x$-axis labels) by different weighting schemes: PNS (weights $w_s$), HH94 [1] and GSC94 [2]. Here, differently from Figure 2, branch lengths in the tree are rescaled by **A** 0.2, and **B** 5. Weights from each scheme are normalized so that the sum over taxa is 1.

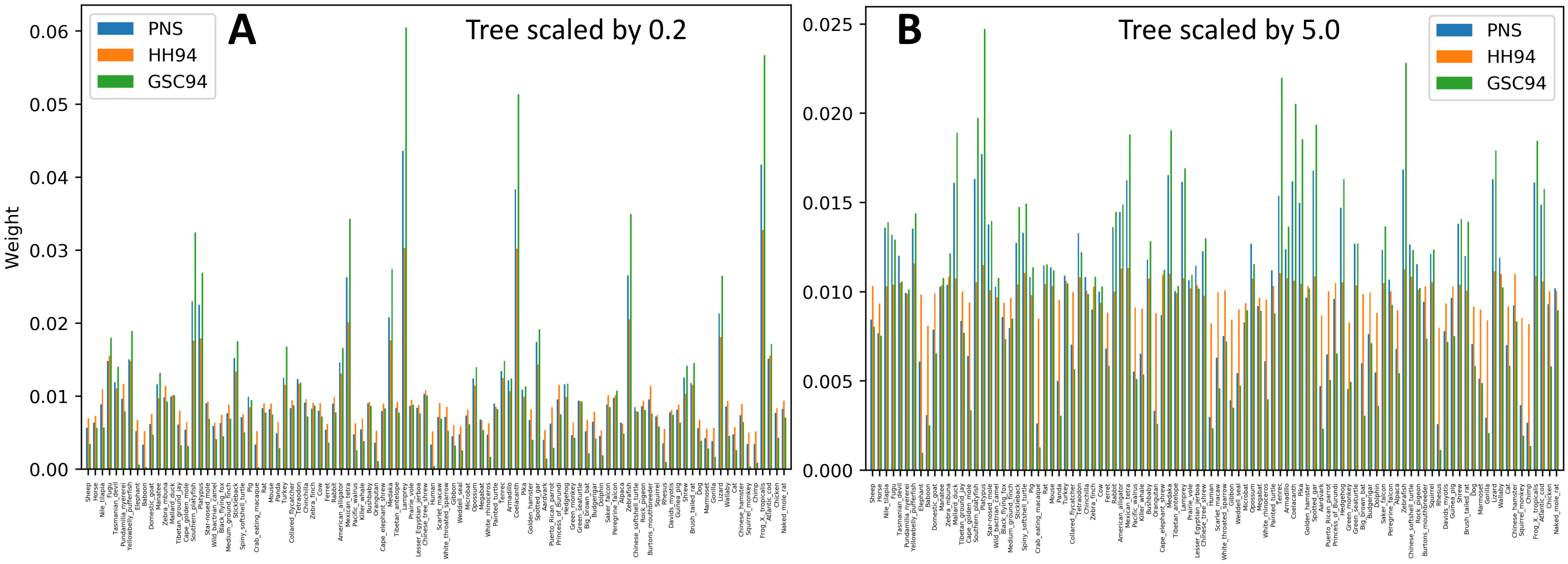

**Figure S3. Computational demand across scenarios.** Violin plots summarise the running times, in seconds, of different methods; each plot contains values for 10 replicates. See Figure 3 for details. Blue plots show running times without accounting for the cost of phylogenetic tree inference, which is instead considered in the orange plots. **A**, **B** and **C** are derived from the simulation scenario with the tree in Figure 1 and nucleotide data. The tree's branch lengths were scaled respectively by (**A**) 0.2, (**B**) 1.0, and (**C**) 5.0. **D** is from the simulation scenario with the tree of Figure 1 and amino acid data. **E** and **F** are from the simulation scenario with the tree in Figure 1 with many human sequences added (100 in **E**, 1000 in **F**) and nucleotide data.

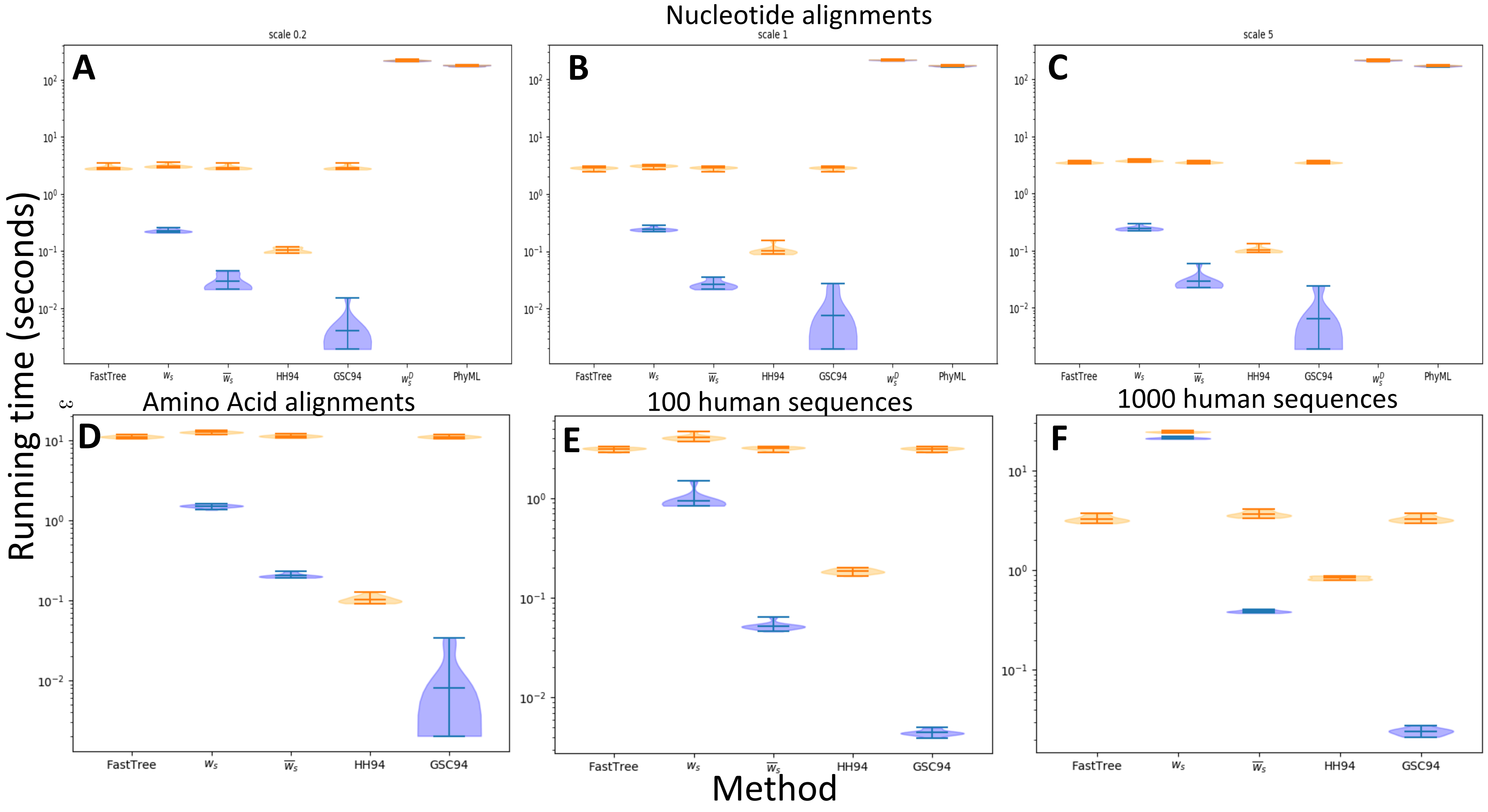

**Figure S4. Equilibrium frequency inference errors of all methods.** Violin plots summarise the character frequency inference error ($y$-axis), measured as the Euclidean distance between the vectors of column-specific simulated character frequencies and inferred ones. Each plot contains 10 replicates, and each replicate contains 800 alignment columns evolved under the background character frequencies (**A**–**C**, **G**–**I**) or 200 alignment columns evolved under equilibrium character frequencies sampled from a Dirichlet distribution with $\alpha = 0.1$ (**D**–**F**, **K**, **L**) and $\alpha = 0.02$ (**J**). Each plot refers to a particular character frequency inference method, indicated on the $x$-axis. '-B' appended to a method name refers to its Bayesian version. '$w_s$ (simu.)' refers to the version of $w_s$ approximated using simulations. In **A**–**G** and **J** simulations are under the tree of Figure 1. In **H** and **K** we consider the tree in Figure 1 with 100 added human sequences (see Methods section). In **I** and **L** we add instead 1000 human sequences. In **G** and **J** we consider amino acid sequences; in all other cases we consider nucleotide sequences.

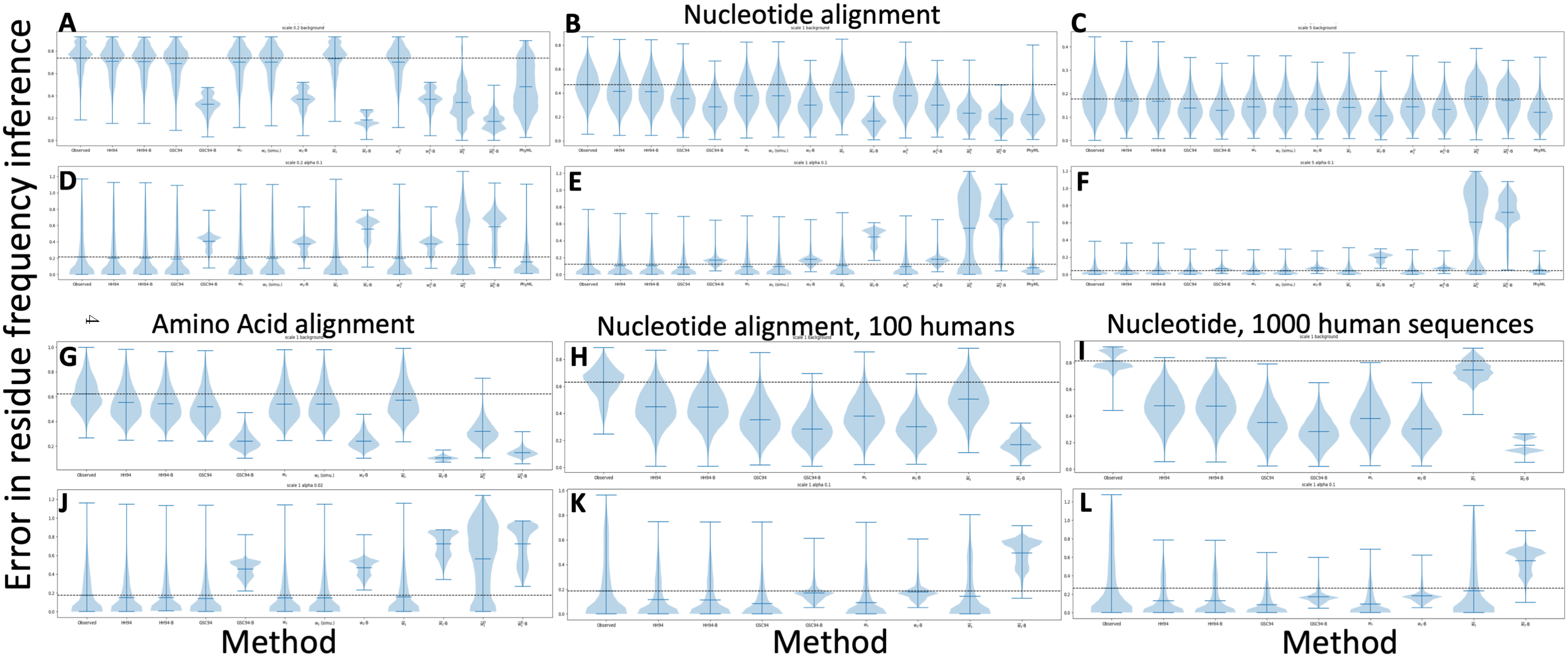

**Figure S5. Conservation score inference error.** **A–F**: Error, measured as the absolute value of the difference between inferred and simulated conservation score (Equation 20). **G–L**: Bias, measured as the difference between inferred and simulated conservation scores. Each plot contains 10 replicates, and each replicate contains 800 alignment columns evolved under the background character frequencies (**A–C**, **G–I**) or 200 alignment columns evolved under equilibrium character frequencies sampled from a Dirichlet distribution with $\alpha = 0.1$ (**D–F**, **J–L**). Each plot refers to a particular method, indicated on the $x$-axis.

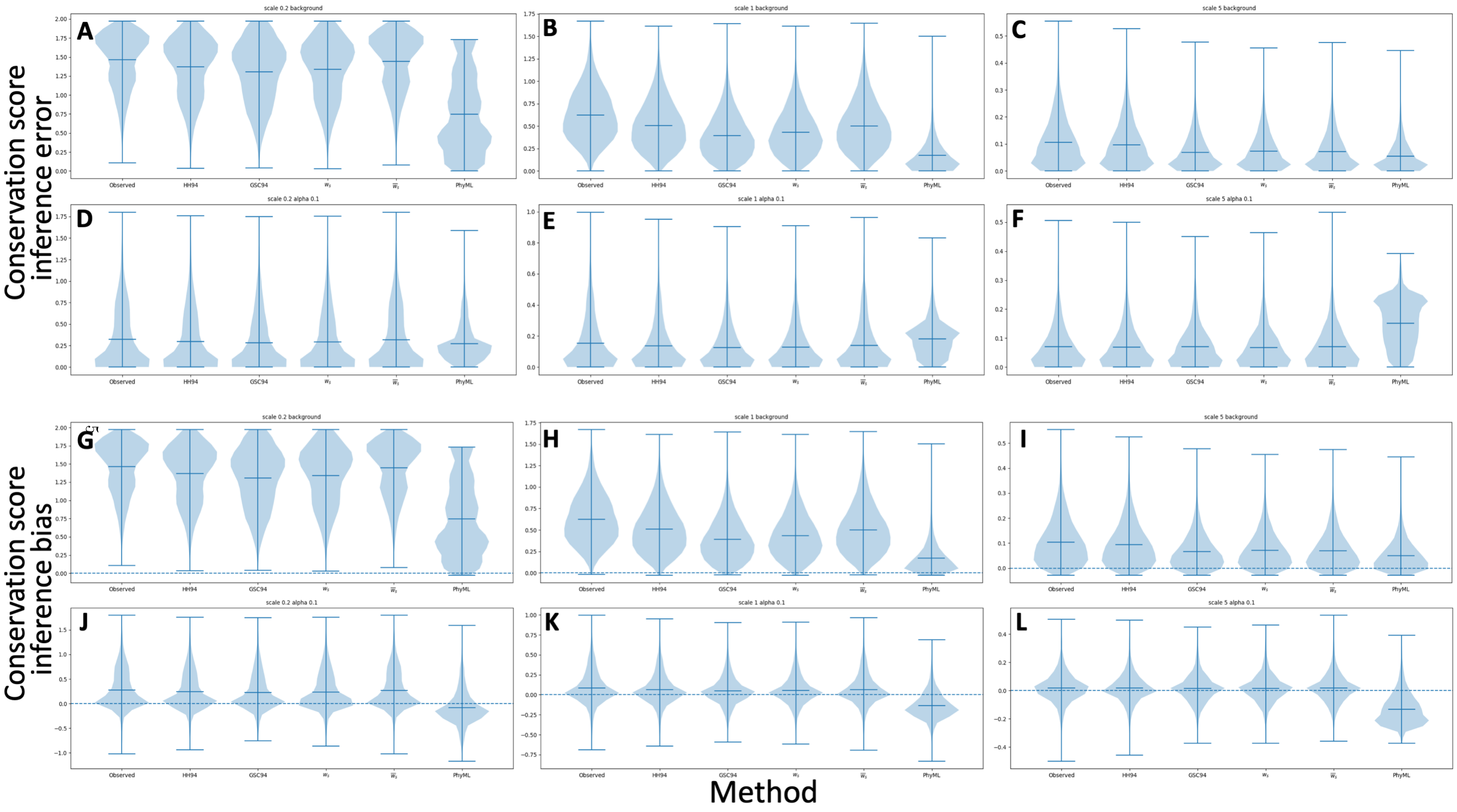

### Supplementary Methods

#### Calculating PNS via simulations I: not conditional on data

It is possible to approximate PNS weights $w_s$ by simulating evolution along a tree. The idea is to repeatedly simulate many alignment columns, and then average (over all simulated columns) the evolutionary novelty of a tree tip to obtain an estimate of its score. At each simulation step, corresponding to one simulated alignment column, we first sample a character at the root of the phylogeny according to the root frequency distribution. We then iteratively sample characters at internal nodes while moving downward on the tree. At each node we condition on the character already sampled at the node's parent. For example, assume that node $\nu_2$ is separated from its parent $\nu_1$ by a branch $b$ of length $t$, and that we have already sampled character $j_1$ at $\nu_1$. We sample a character $j_2$ at $\nu_2$ using probabilities $P_t^{j_1,j_2}$ where $P_t = \exp(tQ)$ is the probability matrix and $Q$ is the substitution rate matrix. If we sample a $j_2 \neq j_1$, then we know that at least one mutation occurred along $b$, and so that no tip descendent from $b$ is PIBD to any tip not descendent from $b$ at this alignment column. There are however two ways in which we can have $j_2 = j_1$: either with no mutation on $b$, with probability $I_t^{j_1}$ (Equation 3), or with more than one mutation on $b$, with probability $P_t^{j_1,j_1} - I_t^{j_1}$. As we move down the tree, we simulate and record which characters are sampled at each node, and whether mutations happened on each branch.

Once we have simulated characters at all tree tips, we add to the partial score $\widehat{w}_s$ of tip $s$ the score corresponding to $1/\mathrm{PIBD}(s)$, the inverse of the number of tips simulated to be PIBD to $s$ at the current alignment column. The final approximation of $w_s$ is obtained, after all alignment columns are simulated, by dividing $\widehat{w}_s$ by the number of simulated alignment columns. In practice, we simulate one block (e.g. 100) of alignment columns at a time, and if the score approximations do not change significantly once a new alignment block has been added (for example the differences between the scores are below a certain threshold $\epsilon$, e.g. $\epsilon = 0.01$), we stop the simulations.

#### Calculating PNS via simulations II: conditional on data

We can also use simulations to approximate $w_s^D$ scores, that is, scores conditional on the data from a specific alignment column. We use a slight modification of the up-down approach of [3] to sample characters at internal nodes of the phylogeny conditional on the observed alignment column $D$. The small difference from [3] is that we do not need to perform the last step of the up-down algorithm, that is, sampling a mutational history within each branch conditional on the two states at the end of the branch. This is because we only need to know if any strictly positive number of mutations (i.e. one or more) happened on a branch, or none. This is done as described in the previous section.

#### Calculating PNS via brute-force

A simple but computationally demanding way to calculate the PNS is to iterate over all possible mutational histories along the phylogeny, calculating the probability of each mutational history and its contribution to the scores. This brute-force method requires exponential time in the number of phylogenetic tips, and we only use it to test the correctness of the other methods and as an example to showcase the properties of the PNS.

In a rooted tree $\phi$ with $N$ tips and $2N-2$ branches, we define a mutational history as a pair of vectors $(\boldsymbol{\mu}, \boldsymbol{o})$. $\boldsymbol{\mu}$ has $2N-2$ boolean entries, and each entry $\mu_b$ is associated with one branch $b$ of the tree. $\boldsymbol{o}$ has $2N-1$ character entries, each entry $o_\nu$ associated with one node $\nu$ of the tree. A value $\mu_b = 0$ represents no mutation happening on $b$; otherwise, $\mu_b = 1$ represents at least one mutation happening on $b$. A value $o_\nu = j$ means that character $j$ is found at node $\nu$ in the mutational history considered. These two vectors are sufficient to describe all the aspects of a mutational history that matter for PNS. However, not all possible vector pairs ($\boldsymbol{\mu}$,$\boldsymbol{o}$) describe a legitimate mutational history. For example, if for a branch $b$ with parent node $\nu_1$ and child node $\nu_2$ we have $\mu_b = 0$, but also $o_{\nu_1} \neq o_{\nu_2}$, then $\boldsymbol{\mu}$ and $\boldsymbol{o}$ are not compatible with each other. For each $\boldsymbol{\mu}$, the number of possible $\boldsymbol{o}$ consistent with it is $B^{1+\sum_b \mu_b}$, with $B$ the number of characters. This follows from the fact that all $\boldsymbol{o}$ consistent with a given $\boldsymbol{\mu}$ can be obtained by first assigning any character to the root, and then a new character on the child node of a branch $b$ that contains mutations ($\mu_b = 1$). We denote by $O_{\boldsymbol{\mu}}$ the space of all $\boldsymbol{o}$ consistent with $\boldsymbol{\mu}$. When conditioning on data $D$ (that is, for weights $w_s^D$), $O_{\boldsymbol{\mu}}$ only contains character vectors $\boldsymbol{o}$ consistent with data $D$.

Given a mutation vector $\boldsymbol{\mu}$, a character vector $\boldsymbol{o} \in O_{\boldsymbol{\mu}}$, and data $D$, the probability $P(\boldsymbol{\mu}, \boldsymbol{o}, D)$ is given by $\pi(o_\rho) \prod_b P(\mu_b, o_{\nu_\mathrm{c}} | o_{\nu_\mathrm{p}})$, where $\rho$ is the root node, $\nu_\mathrm{p}$ and $\nu_\mathrm{c}$ are respectively the parent and child nodes of branch $b$, and $P(\mu_b, o_{\nu_\mathrm{c}} | o_{\nu_\mathrm{p}})$ is the probability of the considered events happening on branch $b$. If the length of $b$ is $t$, we have:

$$P(\mu_b, o_{\nu_\mathrm{c}} | o_{\nu_\mathrm{p}}) = \begin{cases} P_t^{o_{\nu_\mathrm{p}}, o_{\nu_\mathrm{c}}} & \text{if } o_{\nu_\mathrm{p}} \neq o_{\nu_\mathrm{c}} \\ P_t^{o_{\nu_\mathrm{p}}, o_{\nu_\mathrm{p}}} - I_t^{o_{\nu_\mathrm{p}}} & \text{if } \mu_b = 1 \text{ and } o_{\nu_\mathrm{p}} = o_{\nu_\mathrm{c}} \\ I_t^{o_{\nu_\mathrm{p}}} & \text{if } \mu_b = 0 \end{cases} \tag{21}$$

The probability $P(\boldsymbol{\mu}, D)$ is then given by $\sum_{\boldsymbol{o} \in O_{\boldsymbol{\mu}}} P(\boldsymbol{\mu}, \boldsymbol{o}, D)$. Conditioning on $D$, we have $P(\boldsymbol{\mu}|D) = P(\boldsymbol{\mu}, D)/P(D)$, where $P(D) = \sum_{\boldsymbol{\mu}} P(\boldsymbol{\mu}, D)$. For each $\boldsymbol{\mu}$, the corresponding score $S(\boldsymbol{\mu}, s)$ for tip $s$ can be calculated as the inverse of the number of tips that are PIBD to $s$ within $\boldsymbol{\mu}$. We then have $w_s^D = \sum_{\boldsymbol{\mu}} P(\boldsymbol{\mu}|D) S(\boldsymbol{\mu}, s)$; the scores unconditional on data, $w_s$, are obtained in the same way except that in this case $D$ is defined as not containing any information, that is, $O_{\boldsymbol{\mu}}$ contains all $\boldsymbol{o}$ consistent with $\boldsymbol{\mu}$ without any additional restriction, and so also $P(D) = 1$.

#### Calculating ESN via a pruning approach

Recall that the effective sequence number (ESN) is defined as the sum of weights over all tips $s$ of tree $\phi$: $T = \sum_{s\in\phi} w_s$ or $T^D = \sum_{s\in\phi} w_s^D$. If one is only interested in calculating ESN without needing the values of the individual $w_s$ or $w_s^D$, the following fast pruning-like algorithm can be used. The idea is to calculate $T$ for each subtree, starting from the tips and moving upwards; the $T$ at the root $\rho$ is then the final value of interest.

We focus here on weights $w_s^D$ conditional on data $D$. As before, for calculating corresponding values for weights unconditional on data, $w_s$, we simply need to assume $D$ empty (uninformative) in the following. Given alignment column data $D$, we have

$$\begin{aligned} T = \sum_{s\in\phi} w_s^D = \sum_{s\in\phi} \sum_i \frac{p_s^\phi(i|D)}{i} &= \sum_j \pi(j) \sum_{s\in\phi} \sum_i \frac{p_s^\phi(i|D,j)}{i} \\ &= \frac{\sum_j \pi(j) \sum_{s\in\phi} \sum_i \left(p_s^\phi(i,D|j)/i\right)}{P(D)} \end{aligned} \tag{22}$$

where $j$ is any character, $\pi(j)$ is the root frequency of $j$, and $p_s^\phi(i|D,j)$ is the probability that tip $s$ has $i$ PIBD tips in $\phi$ conditional on $D$ and on having character $j$ at the root. Similarly, $p_s^\phi(i,D|j)$ is the probability that $s$ has $i$ PIBD tips in $\phi$ and data $D$, conditional on having character $j$ at the root. We also define scores

$$S^j = \sum_{s\in\phi} \sum_i \frac{p_s^\phi(i,D|j)}{i} \tag{23}$$

and

$$S_\nu^j = \sum_{s\in\phi_\nu} \sum_i \frac{p_s^{\phi_\nu}(i, D_\nu|j)}{i} \tag{24}$$

where now $\phi_\nu$ is the sub-phylogeny comprising the descendants of node $\nu$, and $p_s^{\phi_\nu}(i, D_\nu|j)$ is the probability of having $i$ tips descendent from $\nu$ that are PIBD to $s$, and of data $D_\nu$ at the tips descendent from $\nu$, conditional on having character $j$ in $\nu$. We also define

$$C_\nu^j = \sum_{i>0} p_\nu^{\phi_\nu}(i, D_\nu|j) \ , \tag{25}$$

the probability of $D_\nu$ and of at least one tip descendent from $\nu$ being PIBD to $\nu$, conditional on having character $j$ at $\nu$. For all tips $s$ and characters $j$ we initialise $S_s^j = C_s^j = \delta(j, D_s)$, where $\delta$ is the Kronecker delta.

Given an internal node $\nu$ separated from child $c_1$ by a branch $b_1$ of length $t_1$, and from child $c_2$ by a branch $b_2$ of length $t_2$, and assuming that $S_{c_1}^j$, $S_{c_2}^j$, $C_{c_1}^j$ and $C_{c_2}^j$ have already been calculated, we have that

$$C_\nu^j = I_{t_1}^j C_{c_1}^j P_\nu(D_{c_2}|j) + I_{t_2}^j C_{c_2}^j P_\nu(D_{c_1}|j) - I_{t_1}^j C_{c_1}^j I_{t_2}^j C_{c_2}^j \ , \tag{26}$$

where $P_\nu(D_c|j)$ is the probability of $D_c$ conditional on having $j$ at node $\nu$, which can be obtained recursively with Felsenstein's pruning algorithm.

Similarly,

$$S_\nu^j = \left(\sum_k P_{t_1}^{j,k} S_{c_1}^k\right) P_\nu(D_{c_2}|j) + \left(\sum_k P_{t_2}^{j,k} S_{c_2}^k\right) P_\nu(D_{c_1}|j) - I_{t_1}^j C_{c_1}^j I_{t_2}^j C_{c_2}^j \ . \tag{27}$$

Using both Equations 26 and 27 recursively up the tree, we can calculate $S_\nu^j$ and $C_\nu^j$ for every $\nu$ and $j$ until we reach the the root $\rho$. Once we reach $\rho$, the total ESN score $T$ can be calculated as in Equation 22: $T = \left(\sum_j \pi(j) S_\rho^j\right)/P(D)$. Since Equations 26 and 27 require a constant time (more precisely, proportional to alphabet size for Equation 27), since these steps are performed once for each node, and since the number of nodes is linear in the number of tips, we have that the cost of these steps is $\mathcal{O}(N)$. When conditioning on data, we also need to perform the classical Felsenstein pruning algorithm to calculate $P_\nu(D_c|j)$, but since its cost is linear in $N$ also, the total cost of calculating $T$ with this approach is still $\mathcal{O}(N)$.

#### Calculating fast PNS via a pruning approach

In Equation 14 we introduced the fast PNS, $\overline{w}_s$, which is a fast approximation of the PNS $w_s$. They are defined as $\overline{w}_s = 1/\mathbb{E}[i_\phi(s)] = 1/\sum_{i=1}^N i p_s(i)$, and similarly for their version conditional on observed data, $\overline{w}_s^D$. The fast PNS can be computed very efficiently with an up-down pruning approach, requiring only $\mathcal{O}(N)$ time instead of $\mathcal{O}(N^3)$ as required for the $w_s$. Here we describe such an algorithm. For completeness, we will describe how to calculate $\overline{w}_s^D$; the case for $w_s$ follows by assuming empty (uninformative) data $D$. For a subtree $\phi'$, a node $\nu$, and a character $j$, we will consider the quantities $N(\nu, \phi'|j)$:

$$N(\nu, \phi'|j) = \sum_{i=1}^N i p_\nu^{\phi'}(i, D_{\phi'}|j) \tag{28}$$

and similarly $N(\nu, \phi', j) = \sum_{i=1}^N i p_\nu^{\phi'}(i, D_{\phi'}, j)$. We will also consider the likelihoods for sub-phylogenies (as in the standard pruning algorithm), $L(\nu, \phi'|j) = P(D_{\phi'}|D^\nu = j)$, which is the likelihood of the sub-phylogeny $\phi'$ conditioned on having character $j$ at node $\nu$; similarly, $L(\nu,\phi',j) = P(D_{\phi'}, D^{\nu} = j)$

First, we initialise the conditional expected values and likelihoods at the tips of $\phi$: for every tip $s$ and every character $j$ we set

$$N(s,\phi_s|j) = L(s,\phi_s|j) = \delta(j, D_s) \tag{29}$$

where $\phi_s$ is the phylogeny consisting only of tip $s$.

If we have a branch $b$ of length $t$ separating nodes $\nu_2$ (bottom, or descendant) and $\nu_1$ (top, or ancestral), and if we denote the two subtrees obtained from $\phi$ by removing $b$ by $\phi_1$ and $\phi_2$, with $\phi_1$ containing $\nu_1$ and $\phi_2$ containing $\nu_2$, and assuming we know $N(\nu_2,\phi_2|j)$ and $L(\nu_2,\phi_2|j)$, we can calculate:

$$\begin{aligned} N(\nu_1,\phi_2|j) &= I_t^j N(\nu_1,\phi_1|j) \\ L(\nu_1,\phi_2|j) &= \sum_k P_t^{j,k} L(\nu_2,\phi_2|k)\ . \end{aligned} \tag{30}$$

This means that we can move the expectations and likelihoods ‘up’ on the branches, and we do this starting from the tips.

Given an internal node $\nu$ and the two descendant sub-phylogenies $\phi_1$ and $\phi_2$ it splits $\phi$ into, we can calculate for every character $j$:

$$\begin{aligned} N(\nu,\phi_1\cup\phi_2|j) &= N(\nu,\phi_1|j)L(\nu,\phi_2|j) + N(\nu,\phi_2|j)L(\nu,\phi_1|j) \\ L(\nu,\phi_1\cup\phi_2|j) &= L(\nu,\phi_1|j)L(\nu,\phi_2|j)\ . \end{aligned} \tag{31}$$

Combining the steps of Equations  and  iteratively, we can calculate these expectations and likelihoods for all the internal nodes, starting from the tips and up to the root $\rho$, which concludes the ‘up’ phase. Given the root frequencies $\pi$, and given the two sub-phylogenies of the root $\phi_1$ and $\phi_2$, we can then calculate

$$\begin{aligned} N(\rho,\phi_1,j) &= \pi(j)N(\rho,\phi_1|j) \\ N(\rho,\phi_2,j) &= \pi(j)N(\rho,\phi_2|j) \\ L(\rho,\phi_1,j) &= \pi(j)L(\rho,\phi_1|j) \\ L(\rho,\phi_2,j) &= \pi(j)L(\rho,\phi_2|j)\ . \end{aligned} \tag{32}$$

The second (‘down’) stage of the algorithm proceeds downward on the tree, from the root towards the tips. Again, we assume we have a branch $b$ of length $t$ separating nodes $\nu_2$ (bottom) and $\nu_1$ (top), and we denote the two subtrees obtained from $\phi$ by removing $b$ as $\phi_1$ (containing $\nu_1$) and $\phi_2$ (containing $\nu_2$). This time we assume we know $N(\nu_1,\phi_1,j)$ and $L(\nu_1,\phi_1,j)$, and we calculate:

$$\begin{aligned} N(\nu_2,\phi_1,j) &= N(\nu_1,\phi_1,j)I_t^j \\ L(\nu_2,\phi_1,j) &= \sum_k L(\nu_1,\phi_1,k)P_t^{k,j}\ . \end{aligned} \tag{33}$$

This lets us move downward along a branch. Now, assuming we reach an internal node $\nu$, then given one of its two descendant sub-phylogenies, $\phi_3$, and its top (ancestor) sub-phylogeny $\phi_1$, we can calculate the following for every character $j$:

$$\begin{aligned} N(\nu,\phi_3\cup\phi_1,j) &= N(\nu,\phi_3|j)L(\nu,\phi_1,j) + N(\nu,\phi_1,j)L(\nu,\phi_3|j) \\ L(\nu,\phi_3\cup\phi_1,j) &= L(\nu,\phi_3|j)L(\nu,\phi_1,j)\ . \end{aligned} \tag{34}$$

We apply Equation  twice for each internal node and once for each descendant sub-phylogeny. We keep applying Equations  and  while moving downward in the tree, until we reach the tips. When we reach a tip $s$, after applying Equation  we obtain $N(s, \phi \setminus s, j)$ and $L(s, \phi \setminus s, j)$ for every character $j$. We also have $N(s, \phi_s|j)$ and $L(s, \phi_s|j)$ from the initialisation step. Combining them, we obtain

$$N(s, \phi, j) = N(s, \phi_s|j)L(s, \phi \setminus s, j) + L(s, \phi_s|j)N(s, \phi \setminus s, j) \ , \tag{35}$$

and summing this over all characters $j$ (when $D$ is empty; otherwise we can just take the one observed value $D_s$ for $j$), and normalising by the total likelihood of $D$, $L(\phi)$, we obtain

$$\overline{w}_s^D = \frac{L(\phi)}{N(s, \phi)} = \frac{L(\phi)}{\sum_j N(s, \phi, j)} \ . \tag{36}$$

In total, the algorithm requires using Equations , , and once for each internal node and for each alphabet character; the bottleneck cost is using Equations and , since they have linear cost in the alphabet size $B$ and they need to be used $B$ times for each node. The total cost of this algorithm is therefore linear in $N$, or more precisely $\mathcal{O}(B^2N)$.
